# Modular Splicing is Linked to Evolution in the Synapse-Specificity Molecule Kirrel3

**DOI:** 10.1101/2023.07.25.550563

**Authors:** Dimitri Traenkner, Omar Shennib, Alyssa Johnson, Adam Weinbrom, Matthew R. Taylor, Megan E. Williams

**Affiliations:** Department of Neurobiology, University of Utah, School of Medicine, Salt Lake City, UT 84112, United States

## Abstract

Kirrel3 is a cell-adhesion molecule that instructs the formation of specific synapses during brain development in mouse and Kirrel3 variants may be risk factors for autism and intellectual disabilities in humans. Kirrel3 is predicted to undergo alternative splicing but brain isoforms have not been studied. Here, we present the first in-depth characterization of Kirrel3 isoform diversity in brain using targeted, long-read mRNA sequencing of mouse hippocampus. We identified 19 isoforms with predicted transmembrane and secreted forms and show that even rare isoforms generate detectable protein in the brain. We also analyzed publicly-available long-read mRNA databases from human brain tissue and found 11 Kirrel3 isoforms that, similar to mouse, encode transmembrane and secreted forms. In mice and humans, Kirrel3 diversity arises from alternative, independent use of protein-domain coding exons and alternative early translation-stop signals. Intriguingly, the alternatively spliced exons appear at branch points in the chordate phylogenetic tree, including one exon only found in humans and their closest living relatives, the great apes. Together, these results validate a simple pipeline for analyzing isoform diversity in genes with low expression and suggest that Kirrel3 function is fine-tuned by alternative splicing and may play a role in brain evolution.

**Significance Statement:** Kirrel3 is an important molecule for synapse and circuit formation with gene variants that are associated with neurodevelopmental disorders, yet Kirrel3 function remains largely unknown. Here, we report new isoforms of mouse and human Kirrel3, including secreted and transmembrane forms, that suggest a diverse repertoire of Kirrel3 actions. Importantly, we identified a new Kirrel3 exon only present in humans and the other great apes with potential to play an important role in circuit formation unique to these species.

## INTRODUCTION

Proper wiring of the mammalian brain requires billions of neurons to form synaptic connections with specific neurons. During development, axons extend to correct brain regions and then synapse with suitable neuronal partners, a process largely governed by the cellular environment, morphology, and cell surface proteins. Cell surface proteins help the cell find matching and avoid non-matching synaptic partners (Sanes and Zipursky 2020, Sudhof 2021). They also function in development by contributing to synapse formation and, likely, continue to function throughout adulthood during synapse maintenance, transmission, and plasticity.

The combinatorial expression of synaptic cell surface proteins provides different cell types with a unique identity that can be further tuned by alternative splicing. Alternative splicing of synaptic molecules can affect their protein-protein interactions and intracellular signaling (Sudhof 2017, Ovando-Zambrano, Arias-Montano et al. 2019, Li, Xie et al. 2020, Gomez, Traunmuller et al. 2021, Trotter, Wang et al. 2023). This, in turn, expands the functional repertoire of synaptic genes and adds a layer of regulatory control. Accordingly, splicing factors are important for all aspects of neural development including neuronal health and accurate neuronal wiring (Furlanis and Scheiffele 2018, Traunmuller, Schulz et al. 2023). Despite the emerging importance and widespread use of alternative splicing programs in synaptic proteins (Ray, Cochran et al. 2020), isoform diversity has been characterized for relatively few cell adhesion proteins including Dscam, Neurexin, and the Neurexin binding partners Neuroligin, Teneurin, and Latrophilin (Hattori, Millard et al. 2008, Schreiner, Nguyen et al. 2014, Treutlein, Gokce et al. 2014, Sudhof 2017, Ovando-Zambrano, Arias-Montano et al. 2019, Li, Xie et al. 2020, Gomez, Traunmuller et al. 2021).

Kirrel3 is a single pass transmembrane protein in the immunoglobulin (Ig) superfamily with five extracellular Ig domains and an intracellular PDZ-binding domain. Kirrel3 mediates cell-adhesion and synapse formation via trans-cellular, homophilic interactions and is essential for normal brain connectivity in mice (Prince, Brignall et al. 2013, Martin, Woodruff et al. 2017, Roh, Choi et al. 2017, Brignall, Raja et al. 2018, Taylor, Martin et al. 2020, Wang, Vaddadi et al. 2021). The Kirrel3 gene is composed of many small exons predicted to undergo alternative splicing. So far, three isoforms were reported in skeletal muscle (Durcan, Conradie et al. 2014), but Kirrel3 isoform diversity has not been tested in the brain. Understanding the alternative splicing program of Kirrel3 is expected to provide new insight to its function in synapse and circuit formation. Moreover, due to the fact that human Kirrel3 missense variants are repeatedly identified as risk factors for neurodevelopmental disorders including autism and intellectual disabilities (Bhalla, Luo et al. 2008, De Rubeis, He et al. 2014, Iossifov, O’Roak et al. 2014, Wang, Guo et al. 2016, Yuen, Merico et al. 2016, Li, Wang et al. 2017, Kalsner, Twachtman-Bassett et al. 2018, Guo, Duyzend et al. 2019, Leblond, Cliquet et al. 2019, Hildebrand, Jackson et al. 2020, Taylor, Martin et al. 2020, Zhou, Feliciano et al. 2022), understanding alternative splicing of Kirrel3 may facilitate the study of disease mechanisms that involve this synaptic gene.

Here, we generated a comprehensive list of Kirrel3 isoforms expressed in the mouse hippocampus obtained with targeted long-read mRNA sequencing. In addition, we examined existing long-read transcriptome data from multiple mouse and human brain regions. We identified a total of 19 mouse and 11 human alternative transcripts predicted to encode distinct Kirrel3 proteins including secreted and transmembrane variants. In both mice and humans, Kirrel3 isoform diversity originates in the independent combination of four protein coding exons, two on the extracellular and two on the intracellular side, together with several alternative C-termini. The four alternatively spliced protein coding exons first appear at different critical branching points in the chordate phylogenetic tree. Moreover, we identified a new alternatively spliced protein-coding Kirrel3 exon exclusively present in humans and the great apes (Hominidae), suggesting a key role of Kirrel3 in regulating brain connectivity in these closely related species.

## MATERIALS AND METHODS

### Animals

Kirrel3 knockout mice were described previously (Prince, Brignall et al. 2013) and are backcrossed to the C57Bl/6J strain background. Male and female mice were used in equal numbers and described in the text. All animal experiments were approved and conducted in accordance with [author University] Institutional Animal Care and Use Committee (IACUC).

### Iso-seq sample processing and analysis

RNA was purified from two whole hippocampi per sample and a total of 6 samples. Samples are from two C57Bl/6J P14 males and females, respectively, as well as one P14 male and female knock-out control (KO; (Prince, Brignall et al. 2013). RNA was tested for quality (Agilent TapeStation, RIN ≥ 8.0) and converted to cDNA (NEBNext Single Cell/Low Input RNA Library Prep Kit for Illumina). cDNA from each sample was amplified and barcoded in a mild stringency PCR (20 cycles of 30 seconds at 98°C, 54°C and 72°C), using a target specific forward primer matching exon 2 of mouse Kirrel3 and a universal cDNA-reverse primer (**extended data, Table 1-1**). Barcoded PCR products ≥ 1kb were enriched (ProNex beads) and quantified (TapeStation D5000). Equal amounts of each sample were combined into a single 500 ng SMRTbell library for full-length transcript sequencing on a single SMRTcell using the Sequel II system (Pacific Bio). ISO-seq analysis was performed using R and the R-package BioStrings (Lifschitz, Haeusler et al. 2022). Results were manually confirmed and aligned using ApE plasmid editor software (Davis and Jorgensen 2022). In brief, Iso-seq reads were demultiplexed using unique 16bp barcode sequences and no more than two mismatches as search pattern. This strategy unambiguously identified the sample origin for 85.8% of all reads. Preliminary inspection of Iso-Seq Kirrel3 transcripts was performed by dividing the Kirrel3 gene (NCBI Gene ID: 67703, 550823 bp) into 155 bp fragments, and screening Iso-seq full-length reads for the presence of any of the 3536 resulting fragments. The preliminary inspection revealed one new exon and seven new exon extensions for a total of 29 alternatively spliced genomic segments (**extended data,Table 2-1**). Iso-seq reads with sample IDs were screened for each of these 29 segments with no more than 40% mismatches.

### Sequence Read Archive (SRA) and Phylogenetic Analysis

Published mouse or human PacBio SMRT transcript libraries generated with brain derived tissues or cell lines were screened for Kirrel3 transcripts using the conserved exon 2 sequence and the NCBI blastn suite (Altschul, Gish et al. 1990). Similar to the analysis of mouse transcripts, human Kirrel3 transcripts were first inspected for the presence of any of 3772 elements, each 155 bp in length, that make up the human Kirrel3 gene (NCBI Gene ID: 84623, 584513 bp). The preliminary inspection revealed three new exons and one new exons extension for a total of 27 alternatively spliced genomic segments (**extended data, Table 2-1**). Similar to the Iso-Seq analyses, mouse and human Kirrel3 transcripts were then examined for the presence of the respective alternatively spliced segments (≤ 40% mismatches) and manually aligned using BioStrings and ApE plasmid editor software, respectively. Phylogenetic analyses of mouse exons 6, 18, 19b, and 22 and human exons 3, 16, 17c, 19a, and 21 was performed by comparing nucleotide sequences to published genomes using the blastn suite (Altschul, Gish et al. 1990), direct gene ortholog inspection and by examining relationship of species with matches using the NCBI Taxonomy database (Schoch, Ciufo et al. 2020).

### Fluorescent In-Situ Hybridization Chain Reaction (HCR)

HCR was performed as previously described (Trivedi, Choi et al. 2018). Mouse brains were cryo-sectioned to 30μm slices, mounted on slides, fixed (4% paraformaldehyde) and washed in PBS. Prior to processing samples according to protocol HCR v3.0 (Molecular Probes), slices were treated with 1μg/mL proteinase K-treated (TE-buffer) and equilibrated in SSC buffer. Custom HCR probes were designed and generated by Molecular Instruments based on provided targets (**extended data, Table 4-1**). After nuclear staining with Hoechst in PBS, coverslips were mounted in Fluoromount-G (Southern Biotech Cat# 0100-01) and imaged (Zeiss CLSM 710).

### Immunoblotting

Immunoblotting from mouse brain tissue was conducted by standard techniques. Hippocampi from a 3-week old Kirrel3 wild type or knockout mice were dissected and processed in a Dounce homogenizer in lysis buffer (50mM Tris 7.5pH, 150mM NaCl, 5mM EDTA, 10% Glycerol, 1% CHAPS) with protease inhibitors at 100mg/mL. The lysate was boiled in Laemmli buffer, run in a 10% bis-tris gel, and transferred to a nitrocellulose membrane. The membrane was probed with mouse-anti-Pan-Kirrel3 (Neuromab, RRID: AB_2315857), mouse anti-Kirrel3^exon19b^ (Neuromab, RRID: AB_2341106), and mouse-anti-GAPDH (Millipore, Cat# AB2302, RRID: AB_ 10615768). Secondary antibody was goat-anti-mouse-HRP (Jackson ImmunoResearch, Cat# 115-035-003, RRID: AB_10015289).

### DNA constructs

cDNAs and plasmids were generated using standard PCR-based restriction enzyme cloning (Taylor, Martin et al. 2020). Extracellular FLAG tags were added to mouse Kirrel3 isoforms F (Kirrel3F) and K (Kirrel3K) in frame after the predicted signal-peptide cleavage site (exon 6) and the tagged constructs placed 3’ to mCherry followed by a viral 2A peptide sequence into the mammalian expression vector pBOS (mCherry-2A-FLAG-Kirrel3F/K).

### Cell aggregation assay

CHO cells were transfected with either mCherry-2A-FLAG-Kirrel3F, mCherry-2A-FLAG-Kirrel3K, or pBOS with mCherry only (mCherry-pBOS). After 48 hours, transfected cells were washed and detached (0.01% trypsin) in magnesium-free HEPES buffer (HMF) (137mM NaCl, 5.4mM KCl, 1 mM CaCl2, 0.34 mM Na2HPO4, 5.6 mM glucose, 10mM HEPES, pH 7.4), spun down, and resuspended in HMF. Next, 100,000 cells suspended in 0.5 mL HMF were allowed to aggregate for 90 min (nutator, 37°C) in BSA-coated 24-well plates. For analysis, cells are fixed by adding paraformaldehyde (PFA; 4% final) and allowed to settle over a 24-hour period. All but 0.3 mL of the supernatant are removed, cells and cell aggregates are carefully transferred in the remaining volume to a 96-well glass bottom plate, and the entire well is imaged (Zeiss CLSM 710). Finally, the fraction of aggregated cells in the entire well was determined using ImageJ software.

### Immunocytochemistry and analysis

293T-HEK cells on poly-D-Lysine treated glass coverslips were transfected using PEI following a previously published protocol (Xie, Xinyong et al. 2013). After 24 hours, cells were fixed (4% PFA) and rinsed with PBS and blocked for 30 minutes with 3% BSA/0.1% Triton-X100 in PBS (blocking buffer). Blocking buffer was also used for all subsequent washes and antibody dilutions. For antigen labeling, cells were incubated for 1-2 hours with primary antibody at room temperature, washed 3 times, and incubated for 45 minutes with secondary antibody. After nuclear staining with Hoechst in PBS, coverslips were mounted in Fluoromount-G and imaged (Zeiss CLSM 710). The relative enrichment of membrane proteins in cell-to-cell contacts was measured as ratio of average fluorescence pixel intensity along a cell’s contact versus free membrane using ImageJ software.

### Statistics

For the CHO aggregation experiment, the sample size (n = 3 for all conditions) indicates independent experiments conducted on different days. Groups were compared by a one-way ANOVA followed by pair-wise post-tests. For the cell junctional enrichment assay, we sampled several cells from 3 independent cultures and used a nested one-way ANOVA followed by pair-wise post-tests. In figure 3 we present this data both as a traditional bar graph (showing all data points from each culture as a different color) and with a mean estimation plot.

### Availability of data and materials

Iso-seq sequencing data are available at NCBI under SRA data file number PRJNA992104. Kirrel3 isoform sequence files have been deposited at NCBI. GenBank accession numbers for mouse Kirrel3 mRNA isoforms F-T are OR239801 - OR239815, respectively. All other materials and data that are not commercially available will be freely provided upon reasonable request.

## RESULTS

### A strategy to enrich for full-length Kirrel3 transcripts

Commonly used deep-sequencing technologies rely on short sequence reads (<250bp) that are powerful assets in defining whole cell or tissue transcriptomes (Corchete, Rojas et al. 2020). Short sequence reads are suitable to determine relative transcript frequencies and relative exon usage for all genes expressed. However, short-read sequencing is not suitable to directly determine the complete exon composition of transcripts longer than the 250bp-read limit (Leshkowitz, Kedmi et al. 2022). Recent advances in long-sequence read technologies such as Iso-seq (Kuo, Tseng et al. 2017) or Nanopore sequencing (Wang, Zhao et al. 2021) can generate millions of high-quality reads of >15 kb that allow full-length transcript characterization. To capture full-length Kirrel3 transcripts, we performed Iso-seq experiments on RNA collected from hippocampal tissue from 14-day old C57Bl/6J mice (n = 4, 2 males and 2 females). As a synapse specificity molecule, Kirrel3 is not an abundant transcript nor protein. Based on published RNA sequencing data on mouse brain samples (Bioproject PRJDB7898, 5 replicates), we estimate that Kirrel3 transcripts comprise an extremely small proportion of the overall mRNA content (<0.001%). Therefore, prior to conducting long-read sequencing, we enriched libraries for Kirrel3 transcripts. We accomplished this via a PCR amplification step using a forward primer matching exon 2 of Kirrel3 and a universal reverse primer to a sequence added during the prior cDNA-synthesis step (**Figure 1**). In addition, a unique Iso-seq library barcode was added to each sample in this step to allow for simultaneous sequencing of all samples in a single sequencing run. We chose the target specific primer because exon 2 contains the start codon required for all known and predicted Kirrel3 isoforms. We also conducted iso-seq on samples from 14-day old male and female Kirrel3 knock-out mice (Prince, Brignall et al. 2013) as sequencing and data analyses controls. We find that our modified iso-seq strategy using a Kirrel3-specific primer generated data sets with a total of 1,394,974 reads including 11, 328 Kirrel3 reads (0.8%). Though Kirrel3 remains a small part of the total data set, we estimate that Kirrel3 transcripts were enriched over 1600-fold on average compared to whole-transcriptome libraries from brain tissue.

**Figure 1:**
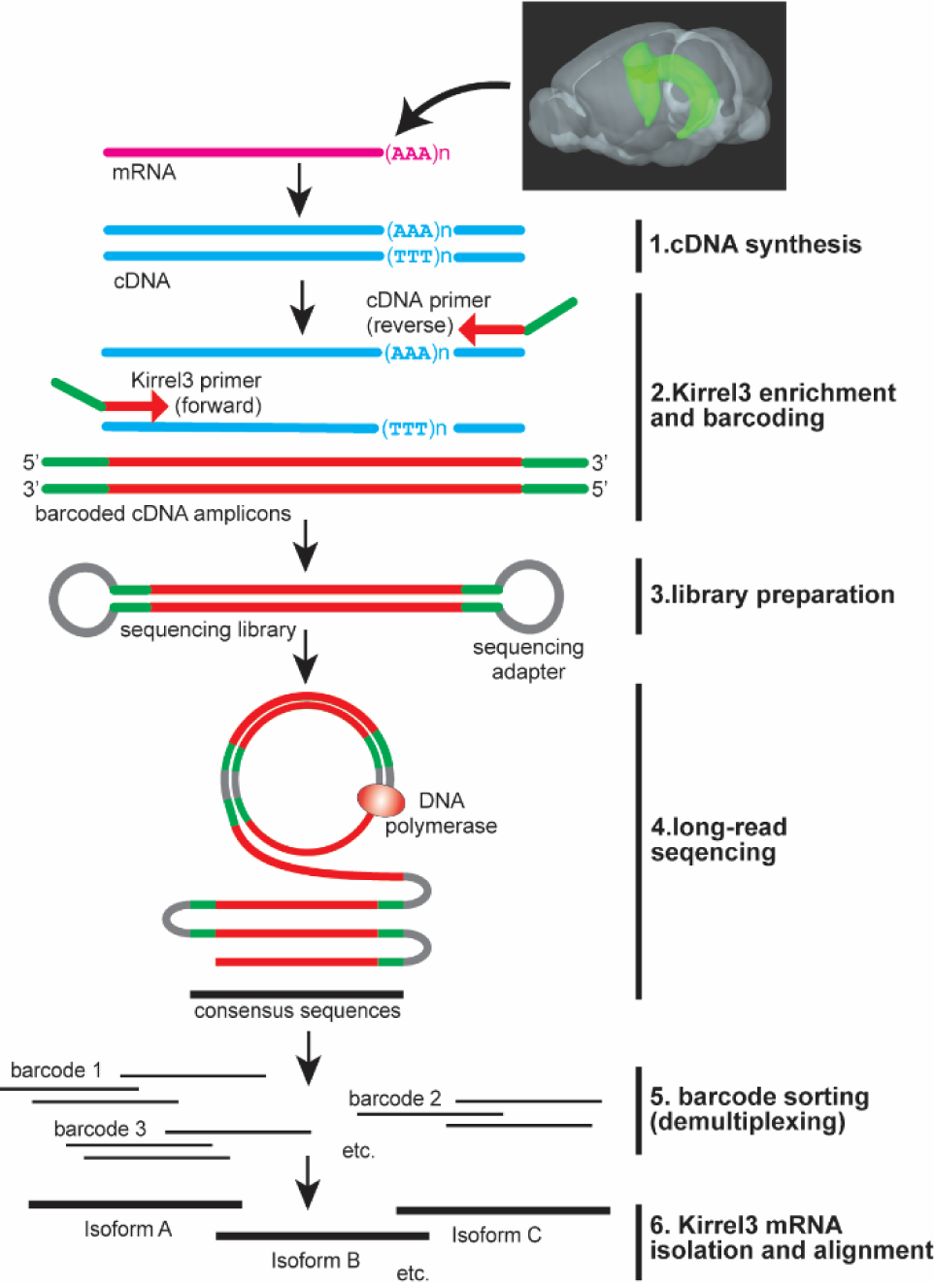
Long-read transcript sequencing. Schematic of the long-read sequencing workflow starting with total mRNA from P14 hippocampal tissue. Pink indicates mRNA, blue cDNA, green DNA-barcodes, red raw Kirrel3 sequences, gray sequencing adapters, and black consensus Kirrel3 sequences. **Extended data Table 1-1** reports the sequence of each barcode used.

### Four independently spliced exons and alternative C-termini generate a diversity of Kirrel3 transcripts

In our custom analysis pipeline, 85.8% of transcript reads could be assigned to a sample based on sample-specific barcodes (**Figure 1 and extended data, Table 1-1**). The remaining 14.2% of all transcripts either contained incomplete or no barcode and were excluded from subsequent analyses. To uncover new Kirrel3 exons, we screened transcripts of each sample for the presence of any known Kirrel3 sequence, including predicted intron sequences (NCBI Gene 67703). Consistent with our PCR strategy to enrich for Kirrel3 transcripts beginning at exon 2, exon 1 of Kirrel3 was absent in our Iso-seq data set. Moreover, hippocampal Kirrel3 transcripts neither contained predicted exons 3 and 4 (ENSEMBL genome browser, ENSMUSG00000032036), nor previously published exon 5, but we included them in the overall gene structure shown in Figure 2A for completeness. Analysis of Kirrel3 transcripts indicates a total of 22 exons, one of which was not previously reported (exon 10 in **Figure 2A**). These exons are alternatively spliced to generate 19 distinct isoforms in mouse hippocampus. Five were previously identified and designated as isoforms A-E. Using the same logic, new isoforms are identified with a letter (**Figures 2B and C**). Based on our data, we also estimated the frequency of each isoform (**Figures 2A and B**) and the cumulative frequency of each predicted protein domain (**Figure 2D**).

**Figure 2:**
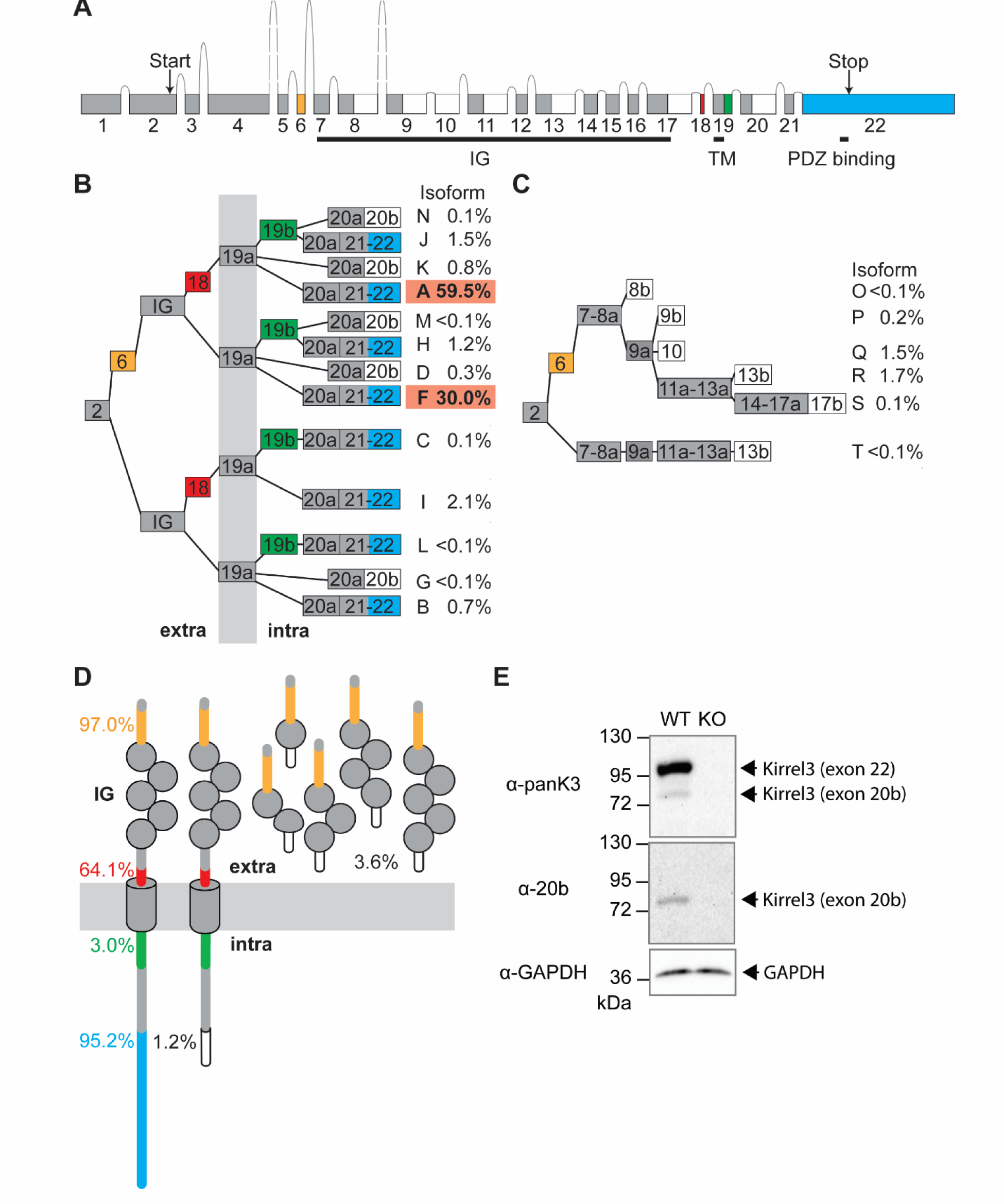
Mouse Kirrel3 gene, transcript isoforms, and proteins. **A)** genomic organization of the mouse Kirrel3 gene, with exons (boxes) and introns (lines), including four independently spliced protein-coding exons (yellow, red, green, and blue). White boxes mark exons or exon parts with a stop-codon. **B)** Alternative splicing of Kirrel3 exons is predicted to produce 13 different transmembrane isoforms. Isoforms are given letters, following the example of previously identified isoforms A-E. Isoform E was not identified in the hippocampus. Exons 8, 9, 11, 13, 17, 19, and 20 are present in transcripts as short (part a) or extended (parts a + b) versions. White boxes indicate exons or exon parts with stop-codon. Percent indicate the relative contribution of a particular Kirrel3-isoform to the total number of complete Kirrel3-transcripts featuring exon 2 and poly A-tail. Exons encode protein segments that are either extra- (‘extra’), intracellular (‘intra’), or spanning the membrane (gray vertical bar). **C)** Six predicted secreted Kirrel3 isoforms (O-T) only comprise extracellular domains. **D)** Schematic of Kirrel3 protein isoforms. Percent are estimates of how frequent protein domains are present in hippocampal Kirrel3 based on isoform-frequencies. **E)** Western immunoblots from hippocampi from 3-week old wildtype and Kirrel3-knockout mice using antibodies directed against Kirrel3 amino acids 46-524 (α-panK3), the peptide encoded by exon 20b (α-20b), or house-keeping gene Glyceraldehyde 3-phosphate dehydrogenase (α-GAPDH). IG: Ig-domain, TM: transmembrane domain. **Extended Data Table 2-1** reports the exact sequences for each mouse and human exon as well as SRA files.

Kirrel3 alternative exons fall into two main groups. The first group contains exons that are either fully included or skipped (exons 6, 18, 22). Exon 6 contains a signal-peptide cleavage site and is present in most transcripts. If exon 6 is skipped, an alternative signal peptide is predicted. As a consequence, slightly different extracellular N-termini are generated dependent on the presence or absence of exon 6. The second group of exons (8, 9, 11, 13, 17, 19 and 20) are found in transcripts with either extended or non-extended 3’ ends. In figure 2, segments that extend an exon are labeled with the letter ‘b’, the respective non-extended exons are labeled with ‘a’. If included in the transcript, six out of the seven b segments (8b, 9b, 11b, 13b, 17b and 20b) introduce a stop-codon (**Figure 2A-C**, white boxes) and produce an alternative C-terminus in the Kirrel3 protein. Exons 8b, 9b, 11b, 13b, 17b, and the newly identified exon 10 are predicted to generate secreted proteins with different numbers of Ig-domains (**Figure 2D**). In contrast, exon 20b is predicted to generate a transmembrane protein with a short intracellular domain lacking the Kirrel3 PDZ-binding domain. Exon 19b does not produce a protein stop, but codes for 25 additional amino acids just intracellular to the transmembrane domain.

The vast majority of isoforms (95%) contain exon 22 and, thus, encode transmembrane proteins with a C-terminal PDZ-binding domain. In contrast, the alternatively generated transmembrane protein with a short intracellular domain lacking a PDZ-binding domain is found in just over 1% of all transcripts. Together with exons 6, 18 and 19b, exon 22 is one of four alternatively spliced exons that appear to be included or excluded from Kirrel3 transcripts independent from each other.

We confirmed our findings by searching published Iso-seq transcript libraries for mouse hippocampus and other brain regions. We found a total of 2237 Kirrel3 transcripts distributed across 21 published mouse Iso-seq libraries (**Extended data, Table 2-1**) but did not detect additional isoforms. Finally, samples from Kirrel3 knockout mouse controls produced the expected 5’ UTR of exon 2 followed by an eGFP coding sequence, which is consistent with how these germline knockout mice were constructed (Prince, Brignall et al. 2013).

### Different Kirrel3 protein isoforms are found in brain tissue

Next, we tested if we could identify distinct Kirrel3-protein isoforms in brain tissue. We initially focused on the existence of isoforms with different intracellular domains because the predicted proteins can be easily resolved by size on a Western blot. We used a pan-Kirrel3 antibody against the extracellular domain and an isoform specific antibody that selectively recognizes isoforms containing the short intracellular domain caused by inclusion of exon 20b. We examined immunoblots of mouse hippocampal lysates using the pan-Kirrel3 antibody and detected two bands that run at the expected molecular weight for full length proteins with the long and short intracellular domains (**Figure 2E**) in wildtype but not Kirrel3 KO tissue. We then blotted the same lysates using the antibody specific to the short intracellular domain only and observed a single band at the lower molecular weight. The results of the immunoblot demonstrate that distinct Kirrel3-isoforms can be identified in brain tissue and that both exon 20b and exon 22 containing mRNA isoforms give rise to proteins in the brain.

### Exons 18, 20b and 22 do not directly affect Kirrel3-mediated homophilic cell adhesion

Homophilic, transcellular binding is necessary for Kirrel3-mediated synapse formation (Taylor, Martin et al. 2020) and structural studies indicate that Kirrel3-Kirrel3 trans binding is mediated predominantly by its first, most N-terminal Ig-domain (Wang, Vaddadi et al. 2021). However, accumulating evidence suggests that intracellular domains can allosterically alter the ligand-binding properties of extracellular domains, either directly by inducing conformation changes, indirectly via additional factors, or by altering the stoichiometric assembly of transmembrane receptors (Changeux and Christopoulos 2017, Ortiz Zacarias, Lenselink et al. 2018, Wang, Heinz et al. 2018, Lara, Burgos et al. 2019). Moreover, a disease-associated missense variant in the intracellular domain was recently identified that significantly attenuates Kirrel3 trans-cellular binding (Taylor, Martin et al. 2020). Thus, we wondered if the C-terminus choice of Kirrel3 affects homophilic cell-adhesion using an established in vitro cell aggregation assay. To test this, we generated expression plasmids encoding cDNAs for isoforms that contain exon 22 (Isoform F) or 20b (Isoform K) (**Figure 2B**). By comparing isoforms F and K, we sought to also obtain information about whether the selectively spliced extracellular domain encoded by exon 18 affects the homophilic transcellular binding of Kirrel3. Exon 18 is present in isoform K, but not in isoform F. We found that suspended cells expressing either Kirrel3 isoform F or K readily form aggregates, but control cells expressing mCherry do not (**Figures 3A, B, Extended Data Table 3-1**). Similarly, both Kirrel3 isoforms are significantly enriched at adherent cell junctions when both contacting cells express Kirrel3 (**Figures 3C-F, Extended Data Table 3-1**). In contrast, neither membrane-GFP nor Neuroligin (a transmembrane protein that does not undergo homophilic binding in trans) are enriched at cell-cell junctions (**Figures 3C-F**). Together, these results show that Kirrel3 isoforms F and K undergo homophilic trans-cellular binding and suggest that the inclusion or exclusion of exons 18, 20b, and 22 does not directly affect the ability of Kirrel3 to bind itself in trans.

**Figure 3:**
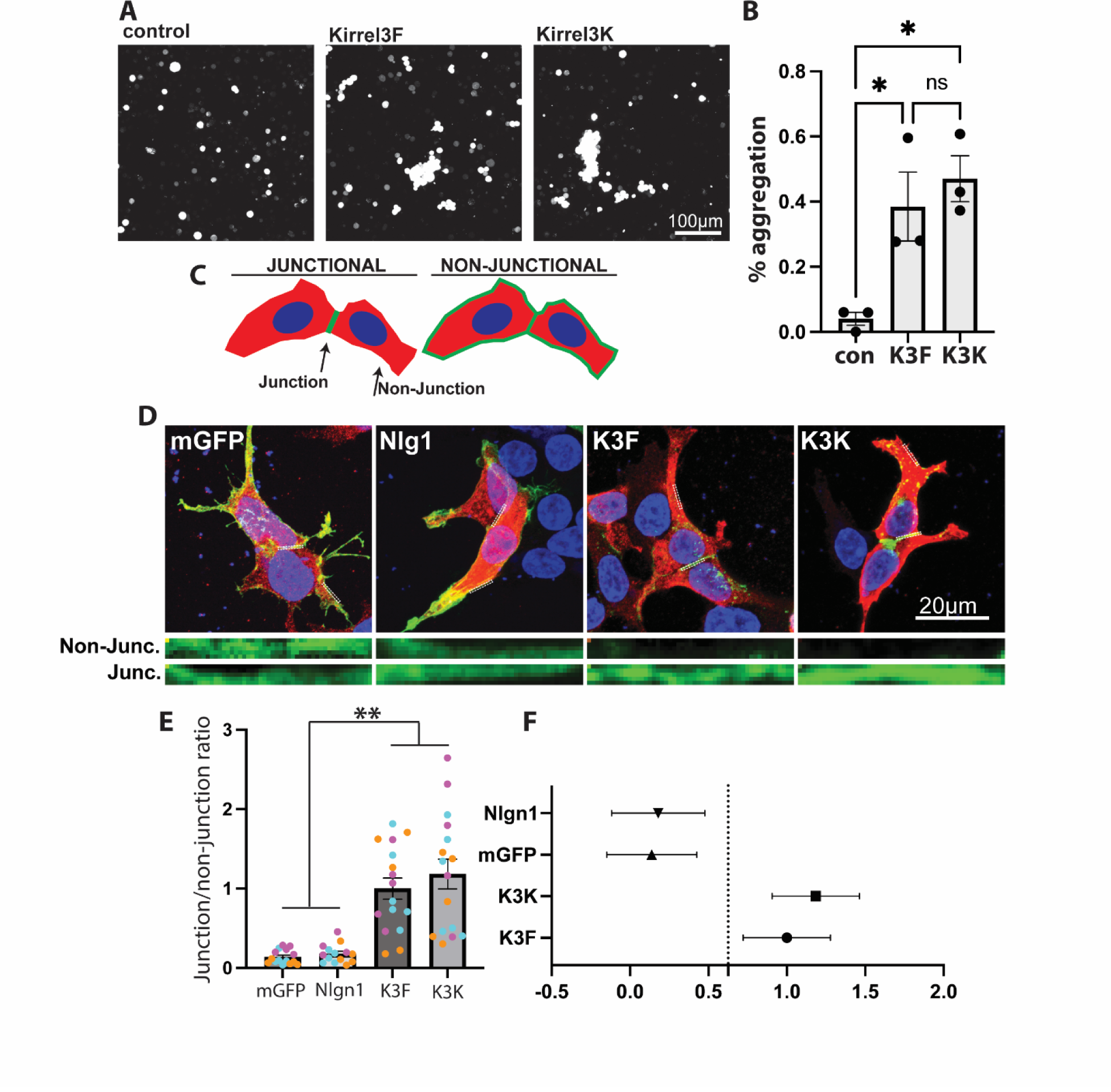
Homophilic binding of Kirrel3 isoforms. **A)** Example of the CHO-cell aggregation assay. CHO cells are expressing mCherry as a control, mCherry-2A-Kirrel3F (K3F), or mCherry-2A-Kirrel3K (K3K). **B)** Quantification of CHO cell aggregation assays. N = 3 independent trials for each condition. Error bars = s.e.m. p=0.01 for a one-way ANOVA and * indicates p<0.05 for each pairwise post-test comparison. **C)** Diagram of the cell junction assay in adherent HEK293 cells. **D)** Example of the cell junction assay. Note that Kirrel3F and Kirrel3K are both highly enriched in the cell-cell junction, but the control proteins, membrane bound GFP (mGFP) and Neuroligin-1 (Nlg1), are not. **E, F)** Quantification of the adherent cell junction assay. **E** shows bar graphs and error bars indicate s.e.m. Each dot indicates a cell and each color denotes cells from an independent culture. One-way nested ANOVA indicates p<0.001 and ** indicates p<0.01 from pairwise post-tests. **F** shows the same data but graphed as a mean estimation plot with error bars showing standard deviation, indicating that Nlg1 and mGFP are significantly different from Kirrel3F and Kirrel3K.

### Kirrel3 isoforms are co-expressed in situ

Next, we sought to test if different cell types or even individual cells express different Kirrel3 isoforms using fluorescent mRNA in situ hybridization (FISH). Because Kirrel3 is not an abundant transcript and most of the alternatively spliced exons are very short, we again focused on the two longest alternatively spliced regions, which encode the alternative intracellular domains exon 22 and 20b. We generated hybridization chain reaction (HCR) FISH probes (Choi, Schwarzkopf et al. 2018) selective to exons 22 and 20b and conducted FISH on brain sections of P14 wildtype and Kirrel3 knockout tissue (**Figure 4 and Extended data, Table 4-1**). Consistent with previous reports (Lein, Hawrylycz et al. 2007, Taylor, Martin et al. 2020, Hisaoka, Komori et al. 2021), we find that Kirrel3 is expressed specifically by DG neurons and GABA neurons in the hippocampus. It is also expressed by many cells in the lateral posterior nucleus (LPN) of the thalamus (**Figure 4**). As predicted by our sequencing results, exon 20b is less abundant than exon 22, but we did not observe obvious differences in the expression pattern of both alternatively spliced exons. FISH probes to smaller exons did not yield any obvious signal. This is likely a technical issue because Kirrel3 transcripts are not highly expressed and the specific probes are very short. Nonetheless, these findings argue against a cell-type specific use of alternative intracellular domains and suggests that most neurons express a mixture of Kirrel3 isoforms with and without a PDZ-binding domain.

**Figure 4:**
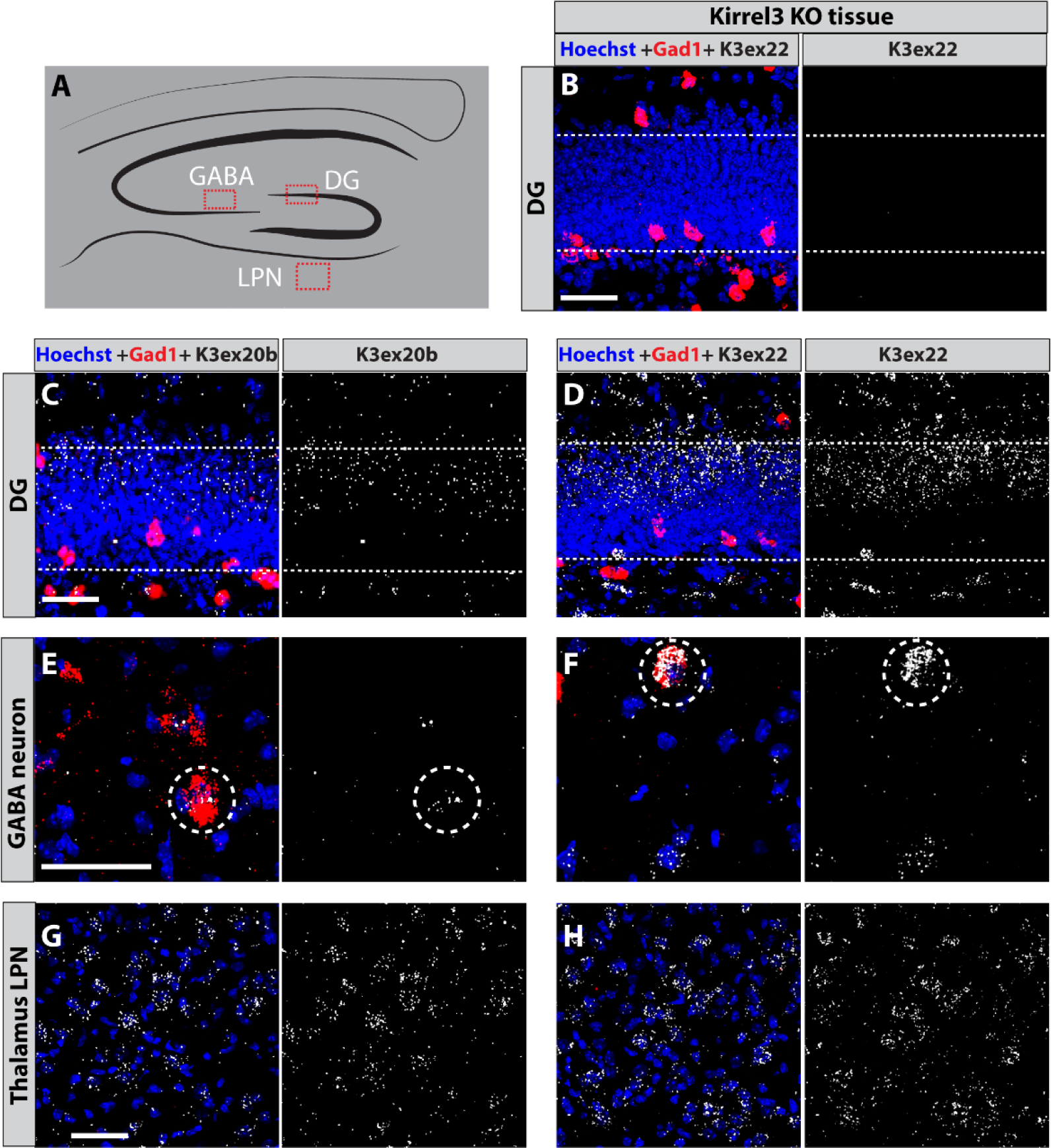
In-situ detection of exons 20b and 22. **A)** Schematic of a coronal section through the hippocampus. Red boxes mark areas imaged in B-H. DG; dentate gyrus, LPN; lateral posterior nucleus of thalamus. **B-H)** Magnified images of regions shown in A that were hybridized with mRNA in situ probes for GAD1 (red) to mark GABAergic neurons, Kirrel3 exon 22 or 20b (white) as indicated to mark specific Kirrel3 isoforms, and the nuclear stain Hoechst (blue). **B)** Kirrel3 knockout tissue produces no Kirrel3 mRNA signal using the larger exon 22 probe. **(C, D)** exon 22 and 20b transcripts are detected in DG neurons of wild type mice. **(E, F)** exon 22 and 20b transcripts are detected in GABAergic neurons (white circles) in area CA3 of wild type mice. **(G, H)** exon 22 and 20b transcripts are detected in the LPN of wild type mice. All scale bars are 50μm. **Extended Data Table 4-1** reports the HCR probe sets used.

### The use of independently spliced Kirrel3 modules is expanded in Hominidae

To examine the extent to which our findings for mouse Kirrel3 isoforms apply to humans, we searched publicly available human brain Iso-seq libraries for reads containing Kirrel3 exon 2 (ENSEMBL genome browser, ENSG00000149571) and identified a total of 1365 transcripts distributed across 19 published libraries. Similar to our strategy for mouse, we screened the identified transcripts for the presence of any Kirrel3 gene sequence (NCBI Gene 84623), including predicted intron sequences. The screen revealed three human Kirrel3 exons and one exon extension not previously reported (**Figure 5A**). The human Kirrel3 gene has 21 exons and 5 alternative C-termini and appears slightly more compact than that of the mouse. Nevertheless, humans have orthologues to the four alternatively spliced mouse Kirrel3 protein coding exons (**Figure 5B**). This includes an alternative signal peptide, insertions just before and after the transmembrane domain, and a short intracellular domain that lacks a PDZ-binding domain. Humans also have Kirrel3 isoforms with a completely new insertion (exon 19a) that adds 30 unique amino acids to the intracellular domain. Intriguingly, across all sequenced genomes a homologue to exon 19a was found only in other great apes (including chimpanzees, orangutans and gorillas), both based on nucleotide and amino acid sequence. This finding suggests an important role of this segment and Kirrel3 in the evolution of the brain. Our search revealed a total of 9 human Kirrel3 transcript isoforms predicted to encode different proteins, including three forms of secreted Kirrel3 (**Figures 5B and C**). The transcript isoforms are designated with a number following the nomenclature of the previously identified isoforms 1-3. Because Kirrel3 transcripts are relatively rare, it is likely that more human isoforms can be found in the future with deeper and targeted sequencing.

**Figure 5:**
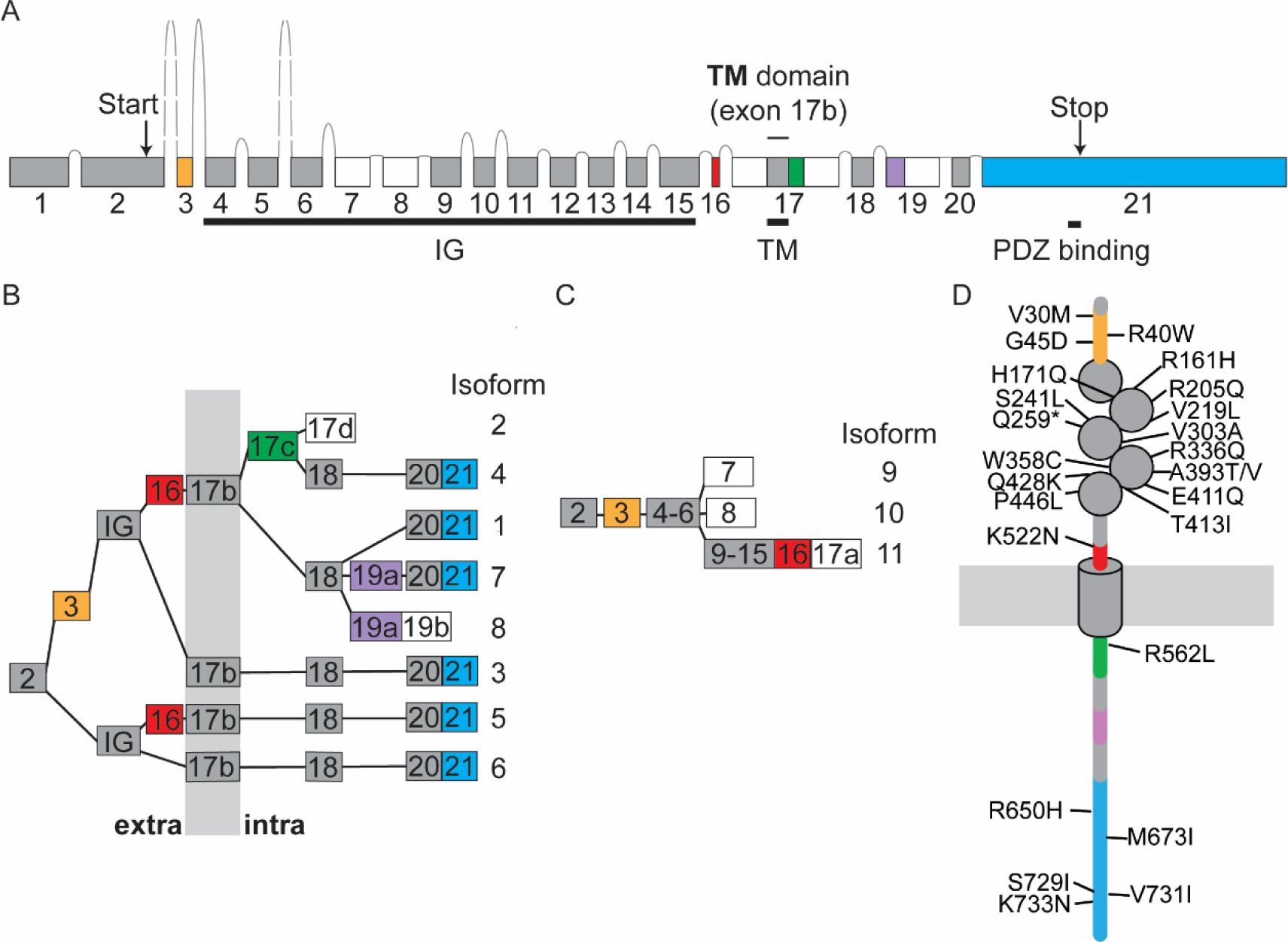
Human Kirrel3 gene, transcript isoforms, and proteins. **A)** Genomic organization of the human Kirrel3 gene, with exons (boxes) and introns (lines), including five independently spliced protein-coding exons (yellow, red, green, purple, and blue). **B)** Alternative splicing of Kirrel3 exons is predicted to produce 8 different transmembrane isoforms. Isoforms are given numbers, following the example of previously identified isoforms 1-3. White boxes indicate exons or exon parts with stop-codon. Exons encode protein segments that are either extra- (‘extra’), intracellular (‘intra’), or spanning the membrane (gray vertical bar). **C)** Three predicted secreted Kirrel3 isoforms (9-11) only comprise extracellular domains. **D)** Schematic of Kirrel3 proteins with the position of protein domains relative to the membrane (horizontal gray bar) and mutations associated with autism. IG: Ig-domain, TM: transmembrane domain.

Kirrel3 variants may be risk factors for autism spectrum disorder and intellectual disabilities in humans and we wondered if identified disease-associated variants are located in specific exons or protein coding domains. We searched the SFARI data base (Banerjee-Basu and Packer 2010, Abrahams, Arking et al. 2013) and the literature for Kirrel3 missense mutations associated with autism or intellectual disabilities (Bhalla, Luo et al. 2008, De Rubeis, He et al. 2014, Iossifov, O’Roak et al. 2014, Wang, Guo et al. 2016, Yuen, Merico et al. 2016, Li, Wang et al. 2017, Kalsner, Twachtman-Bassett et al. 2018, Guo, Duyzend et al. 2019, Leblond, Cliquet et al. 2019, Hildebrand, Jackson et al. 2020, Taylor, Martin et al. 2020, Zhou, Feliciano et al. 2022). Out of the 25 mutations with the strongest predicted association to disease (**Figure 5D**), the majority (15) are within the Kirrel3 Ig-domains 2-5. Interestingly, no mutation with a strong disease link was found in the N-terminal Ig-domain despite its prominent role in homophilic transcellular binding. The remaining mutations are found in one of the alternatively spliced protein modules coded by exon 3 (3 mutations), exons 16 and 17c (1 mutation each), and exon 21 (5 mutations), suggesting important roles for these exons in Kirrel3 function.

### Independently spliced Kirrel3 modules appear at branch points in the chordate phylogenetic tree

Inspired by the identification of an alternatively spliced Kirrel3 exon only present in humans and their closest living relatives, we examined the presence of all 5 alternatively spliced protein coding Kirrel3 segments using both nucleotide and amino acid-based searches across all published genomes. PDZ-binding domain coding exon 22 of mice first appears in combination with the Ig-domains characteristic for Kirrel3 in chordates that evolved over 500 million years ago (Holland 2005) (**Figure 6**). Mouse exon 6, coding for the Kirrel3 N-terminus with a cleavable signal peptide first appears in amniotes, a clade that marks the transition of tetrapods from aquatic to terrestrial habitats about 300 million years ago (Benton and Donoghue 2007). Mouse exons 18 and 19b, both encoding protein segments in juxtaposition to the transmembrane domain, first appear together in placental mammals which evolved in the late cretaceous around 90 million years ago (O’Leary, Bloch et al. 2013). Finally, human exon 19a and its orthologues appear to be the latest addition to Kirrel3 about 12-16 million years ago when the last common ancestor of all great apes lived (Chen and Li 2001).

**Figure 6:**
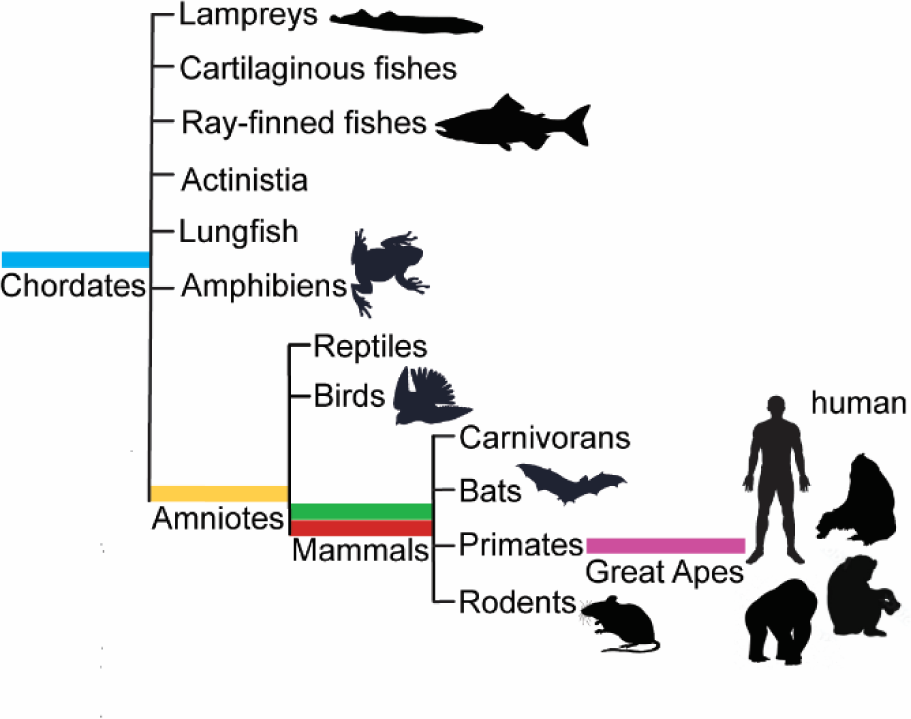
Phylogenetic tree of Kirrel3. The independently spliced protein-coding exons (yellow, red, green, purple, and blue) that produce the Kirrel3 isoform variety first appear at branching points in chordate evolution.

## DISCUSSION

Here, we detect an extensive repertoire of Kirrel3 mRNA isoforms in the mouse brain using a modified long-read sequencing strategy that substantially enriched Kirrel3 transcripts in the dataset. Gene specific enrichment was an essential prerequisite to discover the extensive alternative splicing of Kirrel3, because expression of this target recognition molecule is inherently sparse and is often limited to a subset of cell types within a tissue. Consequently, we find that bulk and single cell sequencing data sets generally contain limited reads of Kirrel3 transcripts. Moreover, sequencing full-length transcripts was necessary because classic next generation sequencing generates only short reads that identify single or few exons, but not complete exon contigs.

Importantly, we provide evidence that even rare Kirrel3 isoforms present in our full-length transcript data set make detectable levels of mRNA and protein in the mouse brain. For example, exon 20b is present in only about 1% of Kirrel3 transcripts, yet we can detect both the corresponding mRNA and protein in brain tissue using a selective in-situ probe and antibody, respectively. Moreover, we demonstrate that exon 20b-containing transcripts give rise to functional proteins. Inclusion of exon 20b produces a Kirrel3 variant characterized by a truncated intracellular domain lacking the PDZ-binding domain typical for Kirrel3 and present in 95% of all isoforms. Despite the truncated tail, Kirrel3 isoforms with exon 20b are still capable of transcellular homophilic binding and can mediate cell-adhesion. The case of Kirrel3 exon 20b illustrates how the study of alternative splicing of synaptic receptors can directly instruct future studies. For example, it would be important to examine how Kirrel3 proteins with truncated tails alter and modulate the function of Kirrel3 isoforms with PDZ-binding domain in forming and maintaining synapses. Addressing this question would also shed new light on how neurons utilize PDZ-domains, a motif present in various scaffolding proteins known to organize and structure synapses.

In a similar vein, it will be important to study the function of secreted Kirrel3 isoforms that, together, constitute an estimated 4% of isoforms. Secreted Kirrel3 isoforms feature one or more Ig-domains and, thus, are predicted to undergo homophilic binding. It is possible that secreted Kirrel3 binds and sequesters transmembrane Kirrel3 variants and prevents their normal function in trans-cellular binding and signaling. Alternatively, secreted Kirrel3 could activate transmembrane Kirrel3 and trigger intracellular signals that are independent of cell-to-cell contact. Another key finding of the present study is that Kirrel3 alternatively uses several protein-coding exons (exons 6, 18, 19, and 22 in mouse) that fall on both the extracellular and intracellular sides of the protein. Other than the C-terminal PDZ-binding domain present in exon 22, the function of the other protein-coding exons remains unknown. However, it is reasonable to assume they are essential for Kirrel3 activity because mutations in the equivalent human domains are associated with disorders. Moreover, these domains first appear at critical branching points in chordate evolution. In this context, the newly discovered intracellular Kirrel3 exon encoding 30 amino acids present only in humans and other great apes is of particular interest as it could serve a synapse-related function unique to Hominidae.

## Supporting information

all supplemental files

