## Supplementary material for "Modular Splicing is Linked to Evolution in the Synapse-Specificity Molecule Kirrel3": all supplemental files

**Extended Data, Table 1-1**

| **Sample** | **Barcode** | **PCR primers** |
|---|---|---|
| wildtype females #1 | ACAGTCGAGCGCTGCG | Forward (5’) |
| wildtype females #2 | ACACTAGATCGCGTGT | Barcode-CTTCTGTGAAAGGAGCCCTTCT |
| wildtype males #1 | CATATATATCAGCTGT | |
| wildtype males #2 | TCACGTGCTCACTGTG | |
| *knockout* females | CACTCGACTCTCGCGT | Reverse (3’) |
| *knockout* males | CACGCACACACGCGCG | Barcode-AAGCAGTGGTATCAACGCAGAGT |

**Table 1. samples and their barcodes.** Each sample represents whole hippocampi of two individuals. Sample cDNA was 5’ and 3’ barcoded using a Kirrel3 specific forward and a universal reverse primer. Wildtype: C567Bl/6J, *KO*: homozygous Kirrel3 knock-out (Prince, Brignall et al. 2013).

**Extended Data, Table 2-1**

### Mouse Kirrel3 gene (NCBI Gene 67703)

| exon | |
|---|---|
| 1 | CTGCTTCCCAAGTTGCTGGCTTTCTGCAATCTCTTGATAGCTACCATTCCCGGTGGCTTTGTGCTCTGAGACTCCGTCCTCTGCGTATCACCCCCTAGATGGTGAGG<br>CTGCCTGCAGGGATGGTGCTCACCCTCCTTGCCCCTCTTGGATGCGCACTACCGGTCTAAGGACTCTCCAAGCTTCTCCATCAGGTGGAGCCGTCCTCTCATCGCCT<br>GCCTCCCTCCTCCCTCGCAGCGAACTGCGAGCAGGCCATCCTACAGAGCCGGGAGGTTAACCGAGCCCCGGAGGAGGTGAGGCCGCCAGCCTGCGCGGGCGG<br>AGATGGCGGGGCCTCCGTCGGGGCCGGCGCCGGCGCCAGAGCCCGGCAGCCTGTGGGTCCTGCAG |
| 2 | CTTCTGTGAAAGGAGCCCTTCTGTCACTCGTCACTCGCTTGCTCGCTCGCCCGCTCGCTAGCTCGTTCGCTCGCTGTGGGAGGAGCCCGCCAGAGGAAGCCGTGT<br>GCCTGGGATGCCAAGAACCAGAGAATGGATCTGCTCCGAGTGGGCACATTGCTAAGGATCCCGGCTTCCCGAGGCGACTGAAAACAAGCATTTGGTTTCGGCTG<br>CCTGCAGATACCCGGAGACACAACGAGACCTAAGCGGACCAGAGGAGGGACAGACCGACTGACAGATAGATGGTCGGCGCGAACCCTGGAGGACCGGCGGCG<br>GAGGCTGAGCACCGCGAGCCCAGCCGCCGCGCTTGAAGAGAAACTAACTGCACACCCAAGTTGCCCGCCGGCTGCCCGCGCGCTGAGGAATGAGACCTTTCCAG<br>CTGGATTTGCTCTTCCTCTGCTTCTTCCTCTTCAGTCAAG |
| 3 | GTAAAGAATAAGAATTCTGGCGGTATACTTCCAGAAGTCGGTAAATGTCAATATTGGATAAGTGTTTCCTTTATATCATGGCAATTTAGCATAAGCTGCTTGGAAA<br>CAAATATCTTCCGCAAGCATACAGTGTTTCA |
| 4 | ACATACCCAGAGCTATGTTTCCATGATGGTTCTACATCCTGTCAAGTGCTCAATCAAGATTAACCATTCTGCCCCTGCAGCACCTGATTCCTTGAAAATCTGCCACC<br>CTCACTAAGCACCAGCCAGCCTCTCAGACACTTCCCTATTCTGGTGAGAATATCTTCAACTCCACTTTGTAGTTGAGGAACTGGAATCTCATTGGGGGTAATTTATT<br>TACCCCACATATAAATAATAGGGGTGAGCCTTCTGTCTCCAGTGGAAATCTCCCCAGCCTTCTGAGCTCTTCTGAGTCAGCCCAGCAGCAGACAGACTCTTACCTTC<br>CCCAGACTCTGAGTCAGGCTTCAGTGCTCCCAGAGAATGAATTTCCTGCAAATCAATCCTGGATGCAGTTATTATTTTATAGGGATGTTGTAAGTTGATAAGGAAG<br>CTGTTATATGCTAATAGCTATTGTAATGTTGTTCTTAAGGCCTATTACTTCTGCTGGTTTGTATTTTAATGTGTGCAAAGGTGATTACAACTTTAATATGAATGTTAA<br>ACAGATTAGCTGATTTATATGTAAAATAAATAAATAAATGGAAACTCATACTCTCA |
| 5 | GACTTCTAAGCCTGTGAAGGATCTTTCCAACTCCCTTCAACTTTTCCAGCTCATTTGTCTTATGCCTCCATGCTGACCTGATGGCTCACCACATGTATTTG |
| 6 | AGCTTGGCCTCCAGAAGAGAGGATGCTGTCTGGTACTGGGCTACATGGCCAAGGACAAGTTTCGGAGAATGAATGAAG |
| 7 | GTCAAGTCTACTCCTTCAGCCAGCAACCCCAGGACCAAGTGGTGGTGTCAGGACAGCCAGTGACTCTGCTGTGTGCCATCCCTGAATATGATGGCTTCGTCCTGT<br>GGATCAAAGATGGCTTGGCTCTGGGTGTAGGCAGAGACCTCTCAA |
| 8a | GTTACCCCCAGTACCTGGTGGTGGGGAACCACCTCTCAGGAGAGCATCACCTGAAGATCCTGAGGGCTGAGCTTCAGGATGATGCCGTGTATGAGTGCCAGGCC<br>ATCCAGGCTGCCATCCGGTCCCGCCCTGCACGCCTCACCGTCCTGG |
| 8b | GTAAGTCACCTACATACAGGTATGCAGGGATGGCATAAGGACTCTGAGAACCTCCAACAGTTCAGCTACTGGAGAGTCCTTCTACAGAGCTTGTCATGGGAGGCT<br>CACCTTTGGATTTCATTGCATCTGGCTCCATCTACTTCTGCCCTCTCCTCTTCCCATCCAAAGGGGACAGGTAAAAGTCCCCCACCTTAGGAAATCATATTGCCACCA<br>ACCTATTATTGTCCTACCTAACCCATGTATTTGGCTGTACGTCGGGAGTAGATGAGGGTCCTAGGTATTCTACAGTGTAAATGAGGCCAGGCCTGCTGCAGCAGT<br>GCCTGTCTGGAAAAGAATGTGACAGAGACTGAAAGACCCTTCTTCCAGAGATGACCTCCAAAAGGATCAGAAAAGCTCTATATGTCCTGACCCCCTCCTAAAAAT<br>GAGTCTCAAGTTTGGTCCTCCATGACTTCAGTGTGCAAAGACCCTTGTTCCAGAGGCCTTCAACCCTGGTCCCCTTGGAGTCCTGGTTCTCAGGACATCCCAGTAA<br>GATTTCATTTCATGTCCTTATTCTTACAATTTTCATTCTTGCTTTTTGTTTCCCATTTTTTATTCAAATGTCACTGGGACAACCTTCCTGAAGAGGTCCCCACTCCTCCA |

| | |
|---|---|
| | TTAAGGGCAGACCTGCCAGAAGATATAGAGAGGTGGGGACTTGTTCAATGACACCTTAAACTTCCCAATTTTAGTTAAGAGTTCGGCCTTGTCACCTCCTTCCTCT<br>GGCCACGTTTTCCAGGGAAGGGGGCTGTTGGAAAACAGTTTGTTCACATACATGATAATCATTCATAATTGC |
| 9a | TGCCACCAGATGACCCCATCATCCTAGGGGGGCCTGTGATCAGCCTTCGGGCAGGGGACCCCCTCAACCTCACCTGCCACGCAGACAATGCCAAGCCTGCGGCTT<br>CCATCATCTGGCTACGTAAAGGAGAGGTCATCAATGGAGCCACCTACTCCAAG |
| 9b | GTGAGCCTGAGGGAGAGAATGGGGGTGCAGGATGGAGAGTGAAACTAAAGATAAGGGGAGACAAGGGTGGTGGTTTCAGGTGCATAGGGACAAGCTCTGCC<br>TCTCTTCTTGTGGAAGTGCTTCTGTTGTTGCTTCACTCCCCCTGTGTGTGCTTGTTTGCTTGCTTGTTGCTTGCTTGCTTGTTGCTTGCTTGCTTGTTTGTTTCATTGCT<br>CACTTTAGTCCTCAGGCCGTTTCTTCCACTACCCTCCCCTTCCCAACCGCCCCATCCTATTCCCCAACTAGAATAACTAGAGGTGAAAACCCTAGAACAAGGTATTTC<br>AGAAGAGGAGACGAGCCAGAAGCAGCCAGGCTAGGATTGCCACAACTAATTCGTGATCCTATGAATAACTGCCTAAATGCCCTGTCTGCCACAAAACCTCCTTAA<br>TTGAATTAAGTCTCAAGAGAGTAAAGATAGAGCCTTTGGAAGGTGGTTATGTCATGAGGGCGTCTACATGAATGAAATTAGTGTCCTTTTAAAAGAGGCCTAAGC<br>ACCAAACCCAGACACCATTGCATATGCCAGCAAGATTTTGCTGAAAGGACCCTGATATAGCTGTCTCCTGTGATGCTATGCCAGTGCCTGGCAAATACAGAAGTG<br>GATGCTCACAGTCATCTATAGGATGGAACACAGGGCCCCCAATGGAGGAGCTAGAGAAAGCATCCAAGGAACTGAAGGGGTCTGCAACCCTATAGGTGGAACA<br>ACAATATGAACTACCCAGTACCCCCAGAGCTCGTGTCTCTAGCTGCATATGTAGCAGAAGATGGCCTAGTCGGTCATCATTGGAAAAAGAGGCCCCTTGGTCTTG<br>CAAACTTTATATGCCCTAGTACAGGGGAACACCAGGCCCAAGAAGTAGGAGTGGGTGGGTAGGGGAGCAGGACAGGGGGTGGGTATAGGGAACTTTTGGGAT<br>AGCATTTGAAATGTAAATAAAGAAAATATCTAATAAAAAATCTATTTGGAAAAAAATTTTAAATAAAATAAAAGAGGCCTAAGAAAGTCCCCTGTCCTTTTCACCA<br>TGTGAAGATGGAAGAGGACAGCCACTAACAAAAAGGCATCTGACCCTTGCCATACACTGGGTCAGTCTGCCCTGGTCTTGGGCTTCTTACCCTCAGGGACTATGA<br>AAAAAAAAAAAA |
| 10 | CTCATCTGGAATCCACAACCTATGCTTCCCTTTTTTCTCTGCCTTGTAGTCCAGAGCAGGATCCTCAGGACCAGCAACCAAAAATATCAGATCTCTCTACAGCCCAG<br>AGCCCACCAAAGCTCATCAAGCTATCCATCCCTAAACCTGTGCAGCCCCTACCCTGCCTCATCCACCCCTTTCCCCACAAACCACAGCAAAGGTTCATCGCACACTC<br>TCTGCTCCACTCCCAGTGGCCTGGTGTGGCATGGCTTACCATGTCCCCTCCTCTTGTGAACTGTGAATTACTACTATGGTCGTGGCTTCTACATGTGTTCATGAATA<br>CAGCCAGAAATAACCAGTGCCACATACCTCCCCAGACGACTTTTATGTCATTCCAGGTGTAGTTTACGTTCTGCTGAGACTTCCAGATGAAGGAAACCTTGGAAGT<br>CCATTGGCCCTTTTTATCTAGGGACAGAAAACTGGGATCTTGAAGGGTGGTGCCTGGACTCAGTTGACTTCTAGCAAACTTAGAAAACCTAGGTTTTCATCTTTTG<br>GGGGGGGGGCTCTCCCAGACAGGCTCAGATCCCTAAGGTCCCCCAAACACCACCAGATGCTATATCAGGATGCCAGGAAAAGCAGCCTCCTGGCTCTGCAGTTG<br>GGCCATAACCTTCCAGGACCCTGACTTGGGGCCCTCAGTTTTAGAGGGTGACAAGGTACATGCTTGTTCTTTAGTTCTGATCCCCAGCTGGCTCCTCCCCATAGGT<br>CCTTAACAGAAGAAATGTCTTTTCCATCAGCAGAGGATGTGAATAAAAGGCCATTCCTATTCTAAGGATGGTTCTGTCCCATCTCAAGTTGGCTCTGCTGCCAAAG<br>GCCACAGATCAAAGCTTGGCGTTACCACCCAGATCCTGGGGCCAGATTCCCAAACAAATGGCTTCTCTTGGGCCTGAATCAACCCCAGGAACAATGCTTTCTCTTG<br>GGAGATAAGTCTGCTGCTCACATCCTGGGTGGAGCCTCACTGTGAGGATGTCAAGAAAGGCCAAGGTCATCCAAGAGAGACTGGGAAGGAGTAGGGTGTTTCT<br>ACATTGGAGGTGCCTGTCCCGGGTCTTTCTGAAGATAGTCCAGCCTGACCCTCCCAGGATACAGAAGAGAAGAGGATACACCTCATCTAGTGAAGACGGGGTCA<br>GGATCACCCTATGAGACCTATTCCAGGGTAGCATACTCAAAAGGCCACTCTCTCCCACACAAGGAAGACAGCTCAGCCAGCCTGGCACATTGCCTTTCTGGAAAG<br>GAGACTGTGTGCCTTTCCTTGCCCAGTTCCACCCTTATTTGGACTGTCACAAGGAGAATGGCATCTGGGCCAGCTGCCAGACCCTGAAGGGCCATGCCAGGCCCG<br>GTTCTCTGTAATACTAGGCAACTGATTTCCTGGGCTCAGTTTCCTCCTCTGAACATGAAAGAGCTGGAAGAAACACTCAGGAAGTGCTCCAGCTGTGACAGGCTG<br>CAAACCCATGGACTGCACCTATCTTAATAGGAGGAGCAGCCTGGCCCAGGAGAACGATAGGACCCAAGCTGAGGGGCTGTCTGAGACTGACCCCCACTCCCCAT<br>CTCCTTCCAGGAGTTCAGTTCAGCCAGGCAGTTTCTTCTCCAGAACTGCAAAGACAAGGGTCATAAGCTAAATAGGACTTCTGACTTACTTACAAAGACAGAGTA<br>GCTGGGACTACACAGTACACACACACACAACCACTCAGACAGGGTCCACAGGAAAGTCTCCAGAACTGGCAGCTTGGGATCTAAGGTCTTCAGGATTTGTCACAT<br>TGTGGTAGTTAAGCACACTGATACTGAAGCTGGGTGGCCTAAATTTGATTCCAGGCTCCACCACCCTCTAGCTATGGGATCTGAACTTAAATTGTCTGTGACCTCG<br>TCTGTATGCTAGGAACATAGTTAGCACCCAGGTCCAAAGATGGCTGGGGCCATGAGCAAGAAACCTGTGTGGCAGGCTGAGCCTGCAGTAAATACAGCTGTTGG<br>GACCATCATGGGTCCTCTGGGTCCTCACACCTGGCCACCTGCATTTTCTGGATGCTGTGGTAGTATCCAAACACAGTAGTCCACAGGGTCCTGCAGGCTGGCCTTC<br>CAGCAAGTCATAATGTTGGCTCAATCTCTCCATCTCAATCTCCCCCAATGAGAAATGGGATTGATAACACCTGTTAGGGCTACCGCAGGACTCAAAAGAAGCCAC |

| | |
|---|---|
| | ACAGCAAAGTTCCTTGCAGAGCTGGTGAGATGACCTTGCATTTGCTGTCGAGCCTGACTACCTTAGCTCCTTCCCCAGGATAGAAAGAGAGAATTAATACCCACA<br>AGTTGTCCTCTGACTCCCACATGCACACTATGACATGTGTGTGCACACA |
| 11a | ACCCTGCTTCGAGACGGCAAACGAGAAAGCATTGTCAGCACCCTCTTCATCTCCCCAGGAGACGTGGAAAATGGACAGAGTATTGTGTGCCGAGCCACCAACAA<br>AGCCATCCCCGGAGGAAAAGAGACCTCTGTCACCATAGACATCCAGC |
| 11b | GTGAGTACCAACTGACCCTCTCTGGCCTGGCCAGGCCTAGGGTTGGGACCAGCCCATGTGGTTGTGTGTGTGTGTGTGTGTGTGTGTGTGTGTGTGTGTATGC<br>AGATTCATTGCAGATTCATACAACCTGTGCATGTGCACAGAGGCCAGAAGAAGTTGTTGGATGTCCTCCTCGATCAACTCTGCCTACTCTTTAGAGGCAACCTCTC<br>TCACTGAACCTGGGTCATATGTTTTCTCTGCTAGGCTAAAGGCCATTAAGTTCAACTGTAAATGTAACTTTTATTTGCCTGGGTGATTATTGTCTCCAAGGCTTACTT<br>CCTCAGTCTGCTATCCTAGGCCTAGTTCTGGAAGCTTCTAGTCTCTGTACAATCTTATCTAGGCCTAGAATGTTTTCAGCCTCTGAGACTTCCTGCTGAATAAGCTCA<br>CCCTTTCTAGTTCTTTTGACTGGCTGATTCAACTCAGTTGTTCTGGATCAAACTCCTCTCCAAACTGACTGATTCAATTACACTTTACTTTCAAATTCTTCTGAATCAT<br>TCTGCTTGTTCTGAAACTAACTCAAGCAATCTGTTCTAATCTGCTAGCTCATTCTCATTCTCGGGCTTGTTCTGTCTTCATCCACGGCTAGCTTGTTCTCCCTCTGCAG<br>CCTGTCTCTGTACAACTAATCCCAGTAAAACTGCCTCCTTCCTCTCTGATGCTCTCTCTTCAAGTAGCTGCTCTTTCTTCTCCTGAGGGTTAGGCACCTCCTATTCTGT<br>CAAATCTTTCTCTGATTCATCACTTTGTCTGCCACTCAATTAGACATTCTGTTCAAACATGGGTACTTCTTTCTACAAACTAACTTTATCTTCATCGTTTGGGATTAAA<br>GGTGTATATAAGGGCACATGTGTATTCCAGCCAGAGGGGTTAAAGGTGTGTGCTAAATGTTGAGCCATACTACAACTAGGAACAGTTTTTTTTTTTTCTGTAAATG<br>ACACAATCTCAGGGTTCACAGTGTGATCAAATATCCTGCAAAACCCCAGCAGTTCTCCTGCCTATGTTCTCCTTGAAGCTGGAGTTATAGTTGTACACCAGGTGCC<br>AGCCTGTTAAGTGGGGGCCAGTATTCAAAATCTCATCCTCATGTTTACGCTGCAAGTGCTTTTAACTACTAAGCCATCTCTCCAGTTCCATAATGCTGATGAATTTA<br>AATACTTTCTAACAGCCCTGGCTCAGTTAGTGCTTCTGGGTTCTGTGTTGTATACACAGCTCTTCGGGGCTAAAGAGTGCACGGGAACTGTGTCTTGGGGTTCTCA<br>GAGCCAGTTTTGACAGCCAGAGGACAGAAGTTTTCTCTGAGCAAATCCTAACTGACACCCTTTAAAATGCAGAAGGAAGAAGAGATGTTTGTTAAGTACTGGGT<br>GAGGCTTAAACACCCAGGCAAACAACTCTCCTCTCCTCTCCTCTCCTCCACTCTCCTCTCCTCTCCTCTCTCTGTATTGCCATTTGTTGTTAACAAACCCCTTCTCCCAT<br>GCCAAGCTTGATCCTCAGTGAGAGTGAGAACAGCAGAATCCTTACCAAGCCTAGGGCTGGGGAGATAAAGAGACTGGAGAGCCAGACCATGGTGGCACACACC<br>TTTAATCCCAGCACTCAGGAGGCAGAGGCAGGAGGATCTCTGTGAGTCTAAGGCCAGCCTGGTCTATAGAATGAGTTTCTAGGACAGCCATGTCTACAGAGAGA<br>AACCCTGTATCAAAAACAAAAACAAAAAGAGACTGGAGAGAAGCACCTCGGCCATGTTCCTCCATCCCTGCCAGGATCCCACCCTGGGTCTTAGCACCATCCAGC<br>AAGCAGATGTGATGAGAGCAAAGCCCAGCTTTGGAAAGTAGGTAAGATGCTAGTTCGGTATGAGGTTCAACATTCCGAACAGATAGATGCATTGCCATGAACTG<br>GGTGGCTCCAGACTTGTCCTTTCACTCTGATAATTCGAATCTCTGGTCTGATTTGCTGATGCTATCCAAGATCTCTTCCTATTCTAGGAGCAATCAAAGCTCTCTCTC<br>AAACACAAGCCAGATCTGGGGGGAATGGGGGTGGAAGACAATAAGTGTCTAAGTCCCTTGCTTCCTGGCTGCCATTTGTGGCTGTGGCACTTCTTAGGCAGGAG<br>GCTATGACAGCTACAGAGTCCCACATAAACATAGTTGGTTCCCCCGAGATGGAAGCCAGGACCTTGACAGAGAAAGATGAAGGAGAAAGAACATCTCTGCCCAA<br>GACTGATATCTGTTGCCCCAAGAGAACCTAGGGACAAGAATGCTATCATGAGAGTATCCCACATACATCCAAATGCATACCCCCTCCAGATGCACTCCACACAAA<br>GGACTGGTTGACTGGCTACATGGTTAACAGGGCTATACCTGAAAGGACCTGGGTATGAAGCTAGCATGAGGCTTAGACCCAGGTTTCAGATTTCCCAGTTGGAA<br>TCTTTCCAAGGCAGTACAGAATAGTGGTTGTGGTCCAACATTCTCTCTAGTTTGAATTCTGGTTTTGCCTTTTACTGATTAGCCTCATTAGTAAATTGATTTGAGCCT<br>TAGTTTCCTCAAGGCGATCATGTGTACCTCTTAAGGTCACAGCTAGAATTAAATGACATCATTCATGTCTATTCTTTGCACCATGCATGACATATTTTGATTAAAGT<br>ATATATTATGGTTGAAACCACGGTGGTGAGTGTGTGGTGACCCTTCCCCAGATGTCATCACATCTGTATCCATTTGATCCAACACTCCATCTCCCCTAGCTGGGAG<br>GTGTGTTTATGTATACCTAACACGTAAGGTCCAACCCTTCTCCATGCCTTCCTCCTCCTCCTCTTCTTCCTCTTCCTCCTCCTCCTCCACCTCTTCCTCCTCCTCTTC<br>CTTTCTTCTCAAATTCACTTGCTCATTCAGTAAGCATTCAGCTGTTTGCAGTGGATACCACAGGCTGGATCAGAGCCGCCTTTGCAAACCTGCTCATTTTCTCAGTCA<br>ACCACAAAGTCTCCAAACTCAGCAAAAGCGCATTTGTTGTTTGTTTCCCTTGGTCTTTATATGTTACCCAACGGAAGACAAAGCCCCCAGATTGAGTAATGGGAGA<br>GAAAGCGCAATCCCCCTCCCCTGACACACACACACACACACACACACACACACACACACACACACACACAAAGCCTCACCACATCTGGACTCTGTGTCCAGCTGCC<br>AAGTTTCAAATGGATGTGCTATAAAATCACTGAGTCATGAAGTTTCTCTCTGCACCCTCCACACTTCTCCTGAAAATGCACATGTGCGCGCACACACACACACACA<br>GGTTGCTGATGGAAACACTAACAGACGCTCAGCTGTAACATCTGTTCAATGTGATCTCAGTTCCACCCTCCTGCAAGTTATGCATCCGACATCTCAAGTGGGATTA<br>AAATCATGTTTAATAGTTGGGTATGACATAAACTATTTATACCTCTGGAACTTGGCAGACTCCTAGAAACTCATATCCTGAGATGTGGCATCCCAGAGCCTGGGGA<br>TAGAGATGCAAATGAAACTGACTGCAGTGGACCCCTCTGGGTCTATTCTCCTTAGGCCCTAAGGACCAAGGGTGTTACAAGGGAAGGAGACGCTGATAACAGGG |

| | |
|---|---|
| | TATACAGGGCATCCAGCAAATGAAGACCCTCGAAGGGTCCCCCAGTAGGTATGTGTTCCTCCTTCTTGCTTCCCAAGAAATCTCACTCCATAAAGAGCACCAAGAA<br>AAGGTTGAGCCTTTGGAAATTTCTCCCTGAGCTCTGTAGTGGGAATTTTGAACTGACACCATCATATCAAGCAAGAGTTGAACTGAGATTACCTATCACCCACCAG<br>GAATGGAGTTTACAGTGTGAGGCTAGCAATGCACAGGGGTCCCAATGGCAATTAGCAAACTCCTCTGGACAAGCTCCTTTACCAGAGAGGATGCTCAGCTTGGC<br>CAGCCTGCCCTGCACAGGACAGTGAGATTGTCAGCCTTTCCTGAGAAGCTGTGGGAGCCACGGGGAAGATTCTGTAGGACTTGCTGGGAGAGTTGCTGGGAGT<br>CTAGGAAGTGAGAAGAGAATCCTGGTGATCCCTTGCCACAGATACCTTAGTAAGCCCTCGAATCTCACTTAAACGGATTCCAATGGTGAAGACCAGATATACCCT<br>GAAAATTGACGACACCCCTAAACAGCAGGAAGTAGCTTAAGAGACACAACACCCTTATTCCCTCTTCACCATTGGTTTCCCCTCTCCTTTTCTCCCTTTCTTTTTATA<br>ATAACAAGAAGGGAGGAATGTTAGCATCAGGACACTCCAAGCCCAGGCCTAACTATGTGATCCTTGACCTGCATTGCCAGGACAATGACCCTCAATCCCACAATC<br>CACTTGGCTCAGTTCACACCCTCACTGGTGTGACATCATTTTCCGTATAAATTAGGGAATCACCCCCTTCTCGCGCTCTCTCTTTCTTCTCTTTCTTCTCTCCCCCCTCT<br>TTCTCTCTTTTTCCTCTTCTGTGCCCTTCTACCTGCTTCACCGCCCCTCTATCTCTGCACAATAAACCTCTCCCATGTGGAACCACCTTG |
| 12 | ATCCACCGCTTGTCAACTTGTCCGTGGAACCACAGCCGGTATTGGAGGACAACATCGTCACGTTCCACTGCTCTGCAAAGGCCAACCCAGCTGTCACCCAGTACA<br>G |
| 13a | GTGGGCCAAACGGGGTCACATCATCAAGGAGGCATCTGGGGAGCTGTATAGGACCACGGTGGACTACACATACTTCTCAGAGCCTGTATCCTGTGAAGTAACCA<br>ATGCCCTGGGCAGCACCAACCTCAGCCGCACAGTGGATGTATACT |
| 13b | GTGAGTGTGGGGGCTCAGGGAACCCTGGGATCAGGCAGTGGGTGGTGTGTGGCTGAGACTATAGCTAGACTCATCACACAAACACAGTGTACATGTTGCGTGT<br>TACAGCATCTCCTAGTACCACTCCTGTCTAGATCCTTGACTCCTCCCTTTGAGGCACCATCTCATAGTGCAGTGACCAGTTACTGAGGACTGGGAGGGCTGACCAG<br>CAGGAATCAAGAAACAGCACTACTGTTCTGGTGCCATGGATCCCAATCTTCTCCCAAGGACCCTACCTCCTAATGCATCCACAATGGGGGTTTCAACATGTAAATC<br>TGAGGGGGTCACATGTACAGTCCATAAGTATATCTTATTTCCCTATTATCAAAATGAGGGAGATGTACTCTCAAGGGCATGCAGGGAGTAAGTATCAGGGTGAG<br>GACTGAAGCCAGACAATAACTAAATCCATCACCGATACACACTCCACTGTCCCACTGAGGGGACACATGGACAGCTTCCTTTCCTCACCAGTGGCCCAACAGATG<br>AAGCCAACCTACTTTTAAGGACTAGCCAGATGGATGT |
| 14 | TCGGTCCTCGAATGACCTCAGAGCCTCAGTCACTGCTGGTAGATCTGGGCTCCGATGCTGTCTTCAGCTGTGCGTGGATCGGCAACCCGTCTCTGACCATCGTGTG<br>GATGAAACGAGGTTCTGGTGTG |
| 15 | GTCCTGAGCAATGAAAAGACCCTAACCCTCAAATCTGTCCGCCAAGAGGATGCTGGGAAGTACGTGTGCCGGGCTGTGGTGCCCCGGGTAGGAGCTGGGGAGA<br>GAGAGGTGACCTTGACTGTCAATG |
| 16 | GACCCCCCATCATCTCCAGCACACAGACCCAGCACGCCCTCCACGGAGAGAAGGGCCAGATCAAATGCTTCATCCGGAGCACACCACCGCCTGACCGAATT |
| 17a | GCCTGGTCCTGGAAGGAGAATGTGCTGGAGTCAGGGACATCAGGGCGCTACACAGTGGAGACGGTGAACACGGAGGAGGGAGTCATCTCCACATTGACCATTA<br>GCAACATTGTGCGTGCTGACTTCCAGACCATATACAACTGTACAGCCTGGAACAGCTTTGGCTCTGACACAGAGATCATCCGACTCAAGGAACAAG |
| 17b | GTGAGGACTGCCAGGGTGCCCCGGGGACCAAGGATTGCCGGCCAGATCTAAGGACGGGCCTCACACTCTTCCTCACTGCAATGCAGCCTCACCTTCTGCTGCCAG<br>GGGCACTGGCCCCTCCAAGGAAACAAAGATACTCCGAGTGATGCCCTCTGGGACTGCTCAGGAAAGGGTGACCACAGAGTGAGCTTGGCTGATGGCAGTGATA<br>AACACCCAATGGTGTCAAACACCCTGGCTTCCACACACTCTGCCTTCCCCTGAGGGATCTGAGCCCGAATGCCCTTGCTAGGTTGCGCTTGGACATACATGCTGAG<br>GAATTAGCCCAAACATGATAAAACAGAAGTCCCAAGAATGAGCCAATGCTAAAGCCAAAGAGTAGCGATGCTTATGTGAGGGAGTGGGGGCTGGGCTCAGACC<br>ACTTGCAACCCCCAGGAGACATCTGGACCCTAGAGGAGACTCTGATTAGTACTGGAGTACAGAGAAGCTGGATTTTGAGAGGTGGAGAAAGAGGAAGGAAGCT<br>TACCTCTCTGTAATCCATCACTATGGCCCCATATGCCAACTGCCCTCACTAGTGACGCTAAGAACTTGGTCTCATGCTAGACATGGTAGCATGTGCTACTCTTAGTT<br>ACACAATAGACTGAGGTAGCAGGATCATTTGAGCCCACACATTCAAGAACAACCTGGGGAACATAGCAAGCCCTTTTCTCAGAACAAACAAGAGATCTCCAGTCT<br>CCATTTAAAGTTCTCCAGCTCAGACTACCAAATGGAGAGTGCCAGAAAGATGACAGCCAGTCCAGAGAAGTTCACTCAGCACATAGTCAGTAGCACTAGGTCCTG<br>GAACGGACTAAAGAGCTTCCCTCTGCCAGCTAGTTCCCAGCCACGGGGTTCAGGGGTGTCCTGGACCTCTGGACCAGAAGTTGGAAGCCCAGCAAGTGGTCCCA<br>TCCAGACCCCACCTCCACCCCCCACCCCTGCAAGGCTCAATAAATGAGGTGGCCGGTGATCACACCTACA |
| 18 | GTTCGGAAATGAAGTCGGGAGCCGGGCTGGAAGCAG |

| | |
|---|---|
| 19a | AGTCTGTACCAATGGCCGTCATCATCGGGGTGGCCGTAGGAGCTGGCGTGGCCTTCCTCGTCCTAATGGCAACCATTGTGGCCTTCTGCTGTGCCCGTTCCCAGA<br>GAA |
| 19b | GTACGGGAGGGAGACCCGGGATCTCAGGGAGGGGGACAGAGAAAAAGGCCAGGCTTAGACTGCCCAGGAGAGCAA |
| 20a | ATCTCAAAGGTGTTGTATCAGCCAAAAATGATATTCGAGTGGAAATTGTGCACAAGGAGCCATCTTCTGGCCGGGAGGCTGAGGACCACACCACCATAAAGCAG<br>CTGATG |
| 20b | GTAAGAGCACAGCCTATGCCCCACTCCATCCTGAGCACACAGACTTCCCGATGCTCTCCATACTGCTGACAGGTAGCACATCCAGGCAAACCCCTCCTGCCCCCAA<br>AGTAAGCCGGGCATGCACTCAGCCAATGAGCATTTATTTGTGCCTCTGTGACGTGATGGGGGAGATTCAAATACAACTTTGATGCAGTTCTTGACCTGGATCCTG<br>ACCACTGAGTAAGGAAAGAACCTGTGCTTGAATTCACATGTGTGGCTGCGTACATGTGAATTCAATGTGATCCTCTTCCTGATCCTCTCTGCAAGAACTGGAATCC<br>TCCCAACTGTCCAGGCAGACTCACATGGGGAGAGGCATGGTATAGATATTCGATGCATTTTTTCTCCTCTTCCCTCTTCACACACACACACACACACATCTGCCAGA<br>AACCCTGGTTTCTTCTTGGGAACTGGATATCCACCAAAAGAAAATGCAAAACTTCTCCAGCAACACCAAACTGCCAACTCCTCTTCCTGATCCTCTCTGCAAGAACC<br>CGAATCCTCCCAACTGTCCAGGCAGAAGTCTAGTCAACTATGCCGGAAGCTCACAGAACACTATATCGTGGTGGCATTATCTGGGAACAGTGACAAGAGGGCAA<br>GGAATAAAGGACAAGGGCTTTTCTGAGCCCCAGTGACCCCATCTATGAAATGGACATGAACACAGTAATATACAGAGCCATCATCAGGATTATAGGGTAACAAC<br>CTGAGTACATACGGGGTACAGTAGAGTTTTCCCTCGACATATGTGCAGAGTTCTGTCCTATAATTCCCAGATACCCAAATCTGAATGTGTACAAGTATAAAATGCT<br>ATATAACATTTATACAGATGCTACCTGCTTCCTCTCATATTCATAAATCATCTCTACTTATGAAGTGTAATGTAAGTGCTGTGTAAACAATTACTGTACCATATTGCT<br>TAAGGATTAATGGCGAGTAAAATATGCACGTTTGATACAGGC |
| 21 | ATGGACCGGGGTGAATTCCAACAAGACTCGGTGCTGAAACAGCTGGAGGTCCTCAAAGAAGAGGAGAAGGAGTTTCAGAACCTGAAG |
| 22 | GACCCCACCAACGGCTACTACAGCGTCAACACCTTCAAAGAACACCATTCAACTCCAACCATCTCCCTGTCCAGCTGCCAGCCAGACCTGCGTCCGACAGGCAAAC<br>AGCGTGTGCCCACAGGCATGTCCTTCACCAACATCTACAGCACCTTGAGCGGCCAGGGCCGCCTCTACGACTATGGACAGAGGTTTGTGCTGGGCATGGGCAGC<br>TCTTCCATTGAGCTTTGTGAGCGGGAGTTTCAGAGGGGCTCCCTCAGCGACAGCAGCTCCTTCCTGGACACGCAGTGTGACAGCAGCGTCAGCAGCAGCGGCAA<br>GCAAGATGGCTACGTGCAGTTTGACAAGGCCAGCAAGGCTTCTGCCTCCTCTTCCCACCATTCCCAGTCCTCTTCCCAGAACTCCGACCCCAGCCGACCCCTGCAG<br>CGGCGGATGCAGACTCACGTCTGAGGACCACGCCCTGTGGTGGGGGATGGGCCAAGGAGGAGGACATGGTACATTCTCGTTCTCCAAGGATTGGGGCTACTTT<br>GCAGAGGACCCTAGAACTGGCCACCTCCGGGGTGGTCTCCGAGCACCTCTGTAAACACCTTCCTTCAAAGCTCTGATCAAGCACAAATCTGGCTCCCAGGTGGGA<br>AATGGAGAGTATGCAGCTGAGCGGATAGTGCTCAGGGCCTCTGTCTCTTGCTCTTCCCTAAAGGTCCCTCAACCACCTTGTCCTCCCATGGGCACTCGTGGCAGCT<br>AGAACTTTGCTTTTATGAAACTGCCGTCCACTTTCCTAGCTCCTCTTGCTGCCCATAAGCCATCCCTGGTGTCTGTATTCCTTGCAGCCTTGAGGAACGGAGGACTT<br>TTTCCCAGCACTGAGCTGCTCCGGAGACCCCAGCCTCCCCACTGTGCATAGCCTATACCGCAGAGGCTGGGGCCTGAGAAATGGCCCTGACCAAAGGAGCATCTG<br>CCTGGGAGTCCGCCCCCACTTTGTTTGGTGTTTGTGTCTGTATTCTTGCAGTTCTGTTCTTGGACTTGATACCTCTGCGCTTGGTGGTGGGACTGGCCTATCAGAGT<br>CTAGTGTCCTCAGAGCTGAGGAAGGGAAAGAGGGAAAATGTGAACTCCTGGAGAACAACTGGCCCAACACACCCTGTGCCAGGCTGTGCAGTTCAGAGCCCTC<br>ACCGTCATCCTCACCCCCTGCCCCGTGTTCTCCCTTCCTTCCCACAGCACAATCGAGCTAATCCGAGGAGTGTGAGAACTCCTCTTGTCAGGGTTTTTTGAACAGTT<br>ACTGAAGCGTGCTTCCTGGGAGATGTGGGTTTGAGGGGGTGCTGAAATCTAGGCTGGAGGATGAGACAGACTCTTTCAGCTGATGACCACAAGGAACAATGAT<br>CCATTCTCCAGTAGATAGGACTCTGTGTGCAAGAGGGACAGTTTTCTTCACCTCTTTCCCATCACTCCCCACTTAAGAATAAACGTTAGGGCCATTACCCCCAACAA<br>A |

**Human Kirrel3 gene (NCBI Gene 84623)**

| exon |
|---|

| | |
|---|---|
| 1 | GCGCCGCCCCCGGACGCGAAGGCTTCCAGCAGGGGCGGCGGTCTCTCTCCTCTCCCCTCTTGCAGCGCACTCCGAGGTCTAGGGACT CTCCGCGCTTCTCCATCAGGTGGAGCCGTCGGCTCCTCGCCGCCTCCCTCCTCCTCCCTCCCCCGGAGCGAACCGCGCGCAGGCAGCC TTGCAGCGCCAGGAGGCTAACCGAGCCCCCGGAGGAGGTGAGGCCGCGGGCAGCCGGGCGGAGATGGCGGGGCCGCCGTCGAGG CTGGCGCCAGAGCCCGACCGGCTGTGGGTCCTGCAG |
| 2 | CTTCAGTGAAAGGAGTCCTTCTGTCACTCGTCACTAGCTCGCTCGCTCACTGTGGGAGGAGCCCGCCTGAGGAAGCCGTGTGCCTGG GATGCCAAGAGCCAGAGAATGGATCTTCTCCGAGTGGGGACATTGCTGACAATCCCGGCTTCCCGAGGCGGCTAAGAACAGGCAGT TTGTGTCGGCTGGCTGCAGATACCCAGAGGCACAAAGAGACCGAAGCCACCCGGAGGGACCCACGGACGGACAGATGGTAGGCGC GAACCCGAGAGGACCGGCGGAGGCTGAGCACCGAGAGCCGCCAAGGAAGAGAAACTAACCACAGCCAAGTTACCCCGCCGGCTTT CCTTCGCTGCGCTAAGGAATGAAACCCTTCCAGCTCGATCTGCTCTTCGTCTGCTTCTTCCTCTTCAGTCAAG |
| 3 | AGCTGGGCCTCCAGAAGAGAGGATGCTGTCTGGTGCTGGGCTACATGGCCAAGGACAAGTTTCGGAGAATGAATGAAG |
| 4 | GCCAAGTCTATTCCTTCAGCCAGCAGCCCCAGGACCAGGTGGTGGTGTCGGGACAGCCAGTGACGCTACTTTGCGCCATCCCCGAAT ACGATGGCTTCGTTCTGTGGATCAAGGACGGCTTGGCTCTGGGTGTGGGCAGGGACCTCTCAA |
| 5 | GTTACCCACAGTACCTGGTGGTAGGGAACCACCTGTCAGGGGAGCACCACCTGAAGATCCTGAGGGCAGAGCTGCAAGACGATGCG GTGTACGAGTGCCAGGCCATCCAGGCCGCCATCCGCTCCCGCCCCGCACGCCTCACAGTCCTGG |
| 6 | TGCCGCCTGATGACCCCGTCATCCTGGGGGGCCCTGTGATCAGCCTGCGTGCGGGGGACCCTCTCAACCTCACCTGCCACGCAGACA ATGCCAAGCCTGCAGCCTCCATCATCTGGTTGCGAAAGGGAGAGGTCATCAATGGGGCCACCTACTCCAAG |
| 7a | TGACAGGATCTTCCTATGTTGCCCAGGCTGGTCACCAAGTCCTAGGCTCAAGCAATCCTCCCACCTCAGCCTCCCAAAGTGCTGGGAT TACAGACATGAGCCTCCACACCCAGCAGACTCTCTTCTCTTCTTCCAAGGACACCAGTCATTGGGTTTAAGGCTCACTCTGAATCCA |
| 7b | GTATGACCTCATCTCAAGAGCCTTAACTAATTACTAATTGCTTAACTAAACTGCAAAGACCCTATTCCCAAATAAGGTCATATTCTGAG GTTCCTGGTAGACAAGAATTTGGAGAGGACACCATTCAACCTGCTGTAGGAGACTAGTACTTATTTCAGGACCAACTTGAAAT |
| 8 | GCCTGAGACTGTCACTCAAGCACCAGAGGCAAAAAGATG |
| 9 | ACCCTGCTTCGGGACGGCAAGCGGGAGAGCATCGTCAGCACCCTCTTCATCTCCCCTGGTGACGTGGAGAATGGCCAGAGCATCGTG TGTCGTGCCACCAACAAAGCCATCCCCGGAGGAAAGGAGACGTCGGTCACCATTGACATCCAGC |
| 10 | ACCCTCCACTGGTCAACCTCTCGGTGGAGCCACAGCCAGTGCTGGAGGACAACGTCGTCACTTTCCACTGCTCTGCAAAGGCCAACCC AGCTGTCACCCAGTACAG |
| 11 | GTGGGCCAAGCGGGGCCAGATCATCAAGGAGGCATCTGGAGAGGTGTACAGGACCACAGTGGACTACACGTACTTCTCAGAGCCCG TCTCCTGTGAGGTGACCAACGCCCTGGGCAGCACCAACCTCAGCCGCACGGTTGACGTCTACT |
| 12 | TTGGGCCCCGGATGACCACAGAACCCCAATCCTTGCTCGTGGATCTGGGCTCTGATGCCATCTTCAGCTGCGCCTGGACCGGCAACCC ATCCCTGACCATCGTCTGGATGAAGCGGGGCTCCGGAGTG |
| 13 | GTCCTGAGCAATGAGAAGACCCTGACCCTCAAATCCGTGCGCCAGGAGGACGCGGGCAAGTACGTGTGCCGGGCTGTGGTGCCCCG TGTGGGAGCCGGGGAGAGAGAGGTGACCCTGACCGTCAATG |

| | |
|---|---|
| 14 | GACCCCCCATCATCTCCAGCACCCAGACCCAGCACGCCCTCCACGGCGAGAAGGGCCAGATCAAGTGCTTCATCCGGAGCACGCCGCCGCCGGACCGCATC |
| 15 | GCCTGGTCCTGGAAGGAGAACGTTCTGGAGTCGGGCACATCGGGGCGCTATACGGTGGAGACCATCAGCACCGAGGAGGGCGTCATCTCCACCCTGACCATCAGCAACATCGTGCGGGCCGACTTCCAGACCATCTACAACTGCACGGCCTGGAACAGCTTCGGCTCCGACACTGAGATCATCCGGCTCAAGGAGCAAG |
| 16 | GTTCGGAAATGAAGTCGGGAGCCGGGCTGGAAGCAG |
| 17a | GAGGTGGCAGTGGAGAAGCTGTAACCCTGCTCCCTGGACCCTGAGACTTGGACACCTGAGGGCCACAACTGATGCCCAGCAGACCTGCGGGGAAGCTACTTAGACCTGCAAACACTAAGAGCCGGATGTCCTCTATATTCACCCGAGAAAGGGAAGGGAAGGAGAGAACATACGGAACAAACAAGGGCAGTGACCATAGTCATTTTTCAAAAAAAGCCAGGGCCTATGACTCACTGGCCGCTGTGTGATAGAGCATTCCTGTTTAATGCCAGAGAGAGCTCCATCGTGCCAGCCCCACCCCTCATCTCCGGTCTTCCTGTCCCCTGGCCCTCCCTTCTCCGTGTCTGCCTGCCTCGCTCTCCTTCCTCTGCCCTTTACCTCAGTTCCCCTGCCCTCTCCTCCACCCACCAAGCCCCTGTGTCCTCCACCCTGGTCTTTGCAGTCCCAGCTAGAGCTGACCCCAGTTGGACTCGGCCTGTGCCTGCCCAGGCCCCACTGAGGCATTTCCCAGGAGGGCCCCATGTGGCAGGGGCTGCCCCCGCAGGAACGGGGTGAGATCATGGGGGGATAGTAGGCCATTCCCCATCCGTCATCAGCTGTGCCCGCTTCTCTCCCACAGA |
| 17b | AGTCTGTGCCGATGGCCGTCATCATTGGGGTGGCCGTAGGAGCTGGTGTGGCCTTCCTCGTCCTTATGGCAACCATCGTGGCGTTCTGCTGTGCCCGTTCCCAGAGAA |
| 17c | GTACGGGAGGGAGATCCGGGATCTCAGGGAGGGGGACAGAGAAAAAGGCCAGGCTTAGGCTGCCCCGGAGAGCAA |
| 17d | GTAAGCAGGAGTGCAATGAACAGGGGTCCTAACAGTGCTGTGAGCTCCTGGGGCAGGGAGTGGGTCTGATGCATCGGTGTATGTGAGCCTGGGCAACATGGCGCCTGGCAGAGTGGGCGCTAGGCTGAGGTTGACCTGGACTAGACTGAACTTCATCTGCAGGGCAGCCAGCATTTTGGATTGAACACATAGCTCTTTCAGTCAGGAACTGTACAGAAAGATAGGGGGAAAAGCGGTTTGTGGTTTGATCCTTGCTCTACAAGAGCTGTTAGTCTAGAGAGACCCCATCTCTACAACAAAATAAAAATAAAGAGCTGCTAGTCTCACCAGAAAAGCAGGTCACTCACACAGCTGTGGGGGAGTGGGTGGGGAAGCAATAAAGGAATTGCTTTGAGAAAACTTTT |
| 18 | ATCTCAAAGGTGTTGTGTCAGCCAAAAATGATATCCGAGTGGAAATTGTCCACAAGGAACCAGCCTCTGGTCGGGAGGGTGAGGAGCACTCCACCATCAAGCAGCTGATG |
| 19a | GCAACTGGCCGGCATTTTACAATAAACGTTCAGTCAATGGAATTGAATCAGCCATGGGAGATCTCTGGTGATGCCCTCGCCCAAGCGAGG |
| 19b | AAAGTATGGCCTCGGATACCTGCCCCTCGCCTCCCCTAACTGCCCCTCACCTCACCTACTCACCCACCCCCCTCAACTGCTTCCCTGTCTCCTAGGATGGGGCTCCAGGAACTCATCTTGACTCTTTCACATGGTCCAG |
| 19c | GCAGTGAGGGGTGGTGGGGGTTATACAGGGCAATCTTTTGAGTTTCTGATCTCATCTTTTCCTTGGGGGAAGTGGAGGTTTAGCCACCTCCTATCTACTGAGCAGCCTTTTTTGGGTTGC |

| | |
|---|---|
| 20 | ATGGACCGGGGTGAATTCCAGCAAGACTCAGTCCTGAAACAGCTGGAGGTCCTCAAAGAAGAGGAGAAAGAGTTTCAGAACCTGAAG |
| 21 | GACCCCACCAATGGCTACTACAGCGTCAACACCTTCAAAGAGCACCACTCAACCCCGACCATCTCCCTCTCCAGCTGCCAGCCCGACCTGCGTCCTGCGGGCAAGCAGCGTGTGCCCACAGGCATGTCCTTCACCAACATCTACAGCACCCTGAGCGGCCAGGGCCGCCTCTACGACTACGGGCAGCGGTTTGTGCTGGGCATGGGCAGCTCGTCCATCGAGCTTTGTGAGCGGGAGTTCCAGAGAGGCTCCCTCAGCGACAGCAGCTCCTTCCTGGACACGCAGTGTGACAGCAGCGTCAGCAGCAGCGGCAAGCAGGATGGCTATGTGCAGTTCGACAAGGCCAGCAAGGCTTCTGCTTCCTCCTCCCACCACTCCCAGTCCTCGTCCCAGAACTCTGACCCCAGTCGACCCCTGCAGCGGCGGATGCAGACTCACGTCTAAGGATCACACACCGCGGGTGGGGACGGGCCAGGGAAGAGGTCAGGGCACGTTCTGGTTGTCCAGGGACGAGGGGTACTTTGCAGAGGACACCAGAATTGGCCACTTCCAGGACAGCCTCCCAGCGCCTCTGCCACTGCCTTCCTTCGAAGCTCTGATCAAGCACAAATCTGGGTCCCCAGGTGCTGTGTGCCAGAGGTGGGCGGGTGGGGAGACAGACAGAGGCTGCGGCTGAGTGCGCTGTGCTTAGTGCTGGACACCCGTGTCCCCGGCCCTTTCCTGGAGGCCCCTCTACCACCTGCTCTGCCCACAGGCACAAGTGGCAGCTATAACTCTGCTTTCATGAAACTGCGGTCCACTCTCTGGTCTCTCTGTGGGCTCTACCCCTCGCTGACCAGAAGCTCTACCTACCCCTGTGCCTGTGCTCCCATACAGCCCTGGGGAGAAGGGGATGACGTCTTCCCAGCACTGAGCTGCCCCAGAAACCCCGGCTCCCCACTGCTGCTCATAGCCCATACCCTGGAGGCTGACAAGCCAGAAATGGCCTTGGCTAAAGGAGCCTCTCTCTCACCAGGCTGGCCGGGAGCCCACCCCCAATTTGTTTGGTGTTTTGTGTCCATACTCTTGCAGTTCTGTCCTTGGACTTGATGCCGCTGAACTCTGCGGTGGGACCGGTCCGGTCAGAGCCTGGTGTACTGGGGGGAGGGAGGGAGGAGGGAGCCTGTGCTGACGGAGCACCTCGCCGGGTGTGCCCCTCCTGGGCTGTGTGACCCCAGCCTCCCCACCCACCTCCTGCTTTGTGTACTCCTCCCCTCCCCCTCAGCACAATCGGAGTTCATATAAGAAGTGCGGGAGCTTCTCTGGTCAGGGTTCTCTGAACACTTATGGAGAGAGTGCTTCCTGGGAAGTGTGGCGTTTGAAGGGGCTGGAGGGCAGGTCTTTAAGATGGCGAGACTGCCCTTCTCAGCTGATAAACACAAGAACGGCGATCCTGTCTTCAGTAAGGCTCCACGAGAAGAGAGGAAGTATATCTACACCTCAACCCTCCTAGTCACCACCTGAAATAAATGTTAGGGACACTACTCCA |

**Mouse SRA Kirrel3 transcripts**

| brain region | SRA file | Kirrel3 transcripts | total transcripts |
|---|---|---|---|
| hippocampus | SRX18486306 | 123 | 2,961,269 |
| hippocampus | SRX18486305 | 186 | 3,124,583 |
| hippocampus | SRX18486105 | 147 | 2,100,786 |
| hippocampus | SRX18486104 | 124 | 1,874,595 |

| | | | |
|---|---|---|---|
| hippocampus | SRX18486098 | 95 | 3,038,371 |
| hippocampus | SRX18486306 | 123 | 2,961,269 |
| hippocampus | SRX18486305 | 186 | 3,124,583 |
| hippocampus | SRX18486105 | 147 | 2,100,786 |
| hippocampus | SRX18486104 | 124 | 1,874,595 |
| hippocampus | SRX18486098 | 95 | 3,038,371 |
| hippocampus | SRX18486097 | 100 | 2,991,115 |
| other | SRX1631051 | 22 | 16,225 |
| other | SRX1631053 | 1 | 40,643 |
| other | SRX4218970 | 266 | 5,219,223 |
| other | SRX4515335 | 94 | 1,309,970 |
| other | SRX5989879 | 2 | 169,679 |
| other | SRX5989880 | 10 | 242,895 |
| other | SRX5989882 | 3 | 93,712 |
| other | SRX9178674 | 193 | 10,730,073 |
| other | SRX9178675 | 187 | 9,508,271 |
| other | SRX9178677 | 9 | 3,371,331 |

**Human SRA Kirrel3 transcripts**

| brain region | SRA file | Kirrel3 transcripts | total transcripts |
|---|---|---|---|
| hippocampus | SRX18485725 | 25 | 1,421,315 |
| hippocampus | SRX18485724 | 33 | 1,440,224 |
| hippocampus | SRX9141829 | 64 | 3,176,068 |
| hippocampus | SRX9141830 | 47 | 3,136,724 |
| other | SRX10976497 | 1 | 649,166 |
| other | SRX11141673 | 17 | 1,121,497 |
| other | SRX11141674 | 25 | 1,487,748 |
| other | SRX11141675 | 314 | 20,213,461 |
| other | SRX11141676 | 21 | 4,826,345 |
| other | SRX1743232 | 3 | 1,179,457 |
| other | SRX19335730 | 44 | 2,888,177 |
| other | SRX8068351 | 4 | 6,363,354 |
| other | SRX8068354 | 2 | 28,141,773 |
| other | SRX8068356 | 319 | 184,339,749 |
| other | SRX9014757 | 4 | 355,152 |
| other | SRX9141821 | 2 | 12,994,243 |
| other | SRX9141822 | 13 | 16,258,243 |
| other | SRX9141824 | 226 | 21,071,378 |

| | | | |
|---|---|---|---|
| other | SRX9141825 | 201 | 9,584,363 |

Extended Data Table 3-1

| Figure | Data structure/Normality | Type of test | results |
| --- | --- | --- | --- |
| Aggregation assay (fig 3B) | ND, sample size too small to test | Ordinary one-way ANOVA | P=0.014 |
| Junction assay (fig 3D) | normal distribution | Nested one-way ANOVA | P=0.0005 |

**Table 3-1.** Details of statistical analyses used in Figure 3.

**Extended Data, Table 4-1**

**Mouse Kirrel3 exon 20b probes**

| | |
|---|---|
| 1 | AAGAGCACAGCCTATGCCCCACTCCATCCTGAGCACACAGACTTCCCGATGC |
| 2 | AGTAGAGTTTTCCCTCGACATATGTGCAGAGTTCTGTCCTATAATTCCCAGA |
| 3 | TACCTGCTTCCTCTCATATTCATAAATCATCTCTACTTATGAAGTGTAATGT |
| 4 | TCCATACTGCTGACAGGTAGCACATCCAGGCAAACCCCTCCTGCCCCCAAAG |
| 5 | AGCCGGGCATGCACTCAGCCAATGAGCATTTATTTGTGCCTCTGTGACGTGA |
| 6 | AGTAAGGAAAGAACCTGTGCTTGAATTCACATGTGTGGCTGCGTACATGTGA |
| 7 | TCAATGTGATCCTCTTCCTGATCCTCTCTGCAAGAACTGGAATCCTCCCAAC |
| 8 | CTCCTCTTCCCTCTTCACACACACACACACACACATCTGCCAGAAACCCTGG |
| 9 | AACTGTCCAGGCAGAAGTCTAGTCAACTATGCCGGAAGCTCACAGAACACTA |
| 10 | GGCTTTTCTGAGCCCCAGTGACCCCATCTATGAAATGGACATGAACACAGTA |
| 11 | ATACAGAGCCATCATCAGGATTATAGGGTAACAACCTGAGTACATACGGGGT |
| 12 | GGGGAGATTCAAATACAACTTTGATGCAGTTCTTGACCTGGATCCTGACCAC |
| 13 | TCCAGGCAGACTCACATGGGGAGAGGCATGGTATAGATATTCGATGCATTTT |
| 14 | ACCAAACTGCCAACTCCTCTTCCTGATCCTCTCTGCAAGAACCCGAATCCTC |
| 15 | TCGTGGTGGCATTATCTGGGAACAGTGACAAGAGGGCAAGGAATAAAGGACA |
| 16 | CCCAAATCTGAATGTGTACAAGTATAAAATGCTATATAACATTTATACAGAT |
| 17 | GTGCTGTGTAAACAATTACTGTACCATATTGCTTAAGGATTAATGGCGAGTA |

**Mouse Kirrel3 exon 22 probes**

| | |
|---|---|
| 1 | GACCCCACCAACGGCTACTACAGCGTCAACACCTTCAAAGAACACCATTCAA |
| 2 | CCAACCATCTCCCTGTCCAGCTGCCAGCCAGACCTGCGTCCGACAGGCAAAC |
| 3 | CGTGTGCCCACAGGCATGTCCTTCACCAACATCTACAGCACCTTGAGCGGCC |
| 4 | GGCCGCCTCTACGACTATGGACAGAGGTTTGTGCTGGGCATGGGCAGCTCTT |
| 5 | ATTGAGCTTTGTGAGCGGGAGTTTCAGAGGGGCTCCCTCAGCGACAGCAGCT |
| 6 | TTCCTGGACACGCAGTGTGACAGCAGCGTCAGCAGCAGCGGCAAGCAAGATG |
| 7 | TACGTGCAGTTTGACAAGGCCAGCAAGGCTTCTGCCTCCTCTTCCCACCATT |
| 8 | CAGTCCTCTTCCCAGAACTCCGACCCCAGCCGACCCCTGCAGCGGCGGATGC |

**Mouse GAD1 probes**

| | |
|---|---|
| 1 | ATGGCATCTTCCACTCCTTCGCCTGCAACCTCCTCGAACGCGGGAGCGGATC |
| 2 | AATACTACCAACCTGCGCCCTACAACGTATGATACTTGGTGTGGCGTAGCCC |
| 3 | GGATGCACCAGAAAACTGGGCCTGAAGATCTGTGGCTTCTTACAAAGGACCA |
| 4 | AGCCTGGAAGAGAAGAGTCGTCTTGTGAGCGCCTTCAGGGAGAGGCAGTCCT |
| 5 | AAGAACCTGCTTTCCTGTGAAAACAGTGACCAGGGTGCCCGCTTCCGGCGCA |
| 6 | GAGACCGACTTCTCCAACCTGTTTGCTCAAGATCTGCTTCCAGCTAAGAACG |
| 7 | GAGGAGCAAACTGCGCAGTTCTTGCTGGAAGTGGTAGACATACTCCTCAACT |
| 8 | GTCCGCAAGACATTTGATCGCTCCACCAAGGTTCTGGATTTCCACCACCCAC |
| 9 | CAGTTGCTGGAAGGCATGGAAGGCTTTAATTTGGAGCTGTCTGACCACCCCG |

| | |
|---|---|
| 10 | TCTCTGGAGCAGATCCTGGTTGACTGTAGAGACACCCTGAAGTACGGGGTTC |
| 11 | ACAGGTCACCCTCGATTTTTCAACCAGCTCTCTACTGGTTTGGATATCATTG |
| 12 | ATCGTTGGATGGTCAAATAAAGATGGTGATGGGATATTTTCTCCTGGGGGAG |
| 13 | ATATCCAATATGTACAGCATCATGGCTGCTCGTTACAAGTACTTCCCAGAAG |
| 14 | AAGACAAAAGGCATGGCGGCTGTGCCCAAACTGGTCCTCTTCACCTCAGAAC |
| 15 | AGTCACTATTCCATAAAGAAAGCCGGGGCTGCGCTTGGCTTTGGAACCGACA |
| 16 | GTGATTTTGATAAAGTGCAATGAAAGGGGGAAGATAATTCCGGCTGATTTAG |
| 17 | GCAAAAATTCTTGATGCCAAACAAAAGGGCTATGTTCCCCTTTATGTCAATG |
| 18 | ACCGCAGGCACGACTGTTTACGGAGCATTCGATCCAATCCAGGAAATTGCGG |
| 19 | CTGCTCATGTCCCGGAAGCACCGCCACAAACTCAGCGGCATAGAAAGGGCCA |
| 20 | ATTCTGGTCAAGGAAAAGGGTATACTCCAAGGATGCAACCAGATGTGTGCAG |
